# Linking plankton size spectra and community composition to carbon export and its efficiency

**DOI:** 10.1101/2021.03.08.434455

**Authors:** Camila Serra-Pompei, Ben A. Ward, Jérôme Pinti, André W. Visser, Thomas Kiørboe, Ken H. Andersen

## Abstract

The magnitude and efficiency of particulate carbon export from the ocean surface depends not only on net primary production (NPP) but also on how carbon is consumed, respired, and repackaged by organisms. We contend that several of these processes can be captured by the size spectrum of the plankton community. However, most global models have relatively simple food-web structures that are unable to generate plankton sizespectra. Moreover, the life-cycles of multicellular zooplankton are typically not resolved, restricting the ability of models to represent time-lags that are known to impact carbon export and its efficiency (pe-ratio). Here, we use a global mechanistic size-spectrum model of the marine plankton community to investigate how particulate export and pe-ratio relate to the community size spectrum, community composition, and time-lags between predators and prey. The model generates emergent food-webs with associated size distributions for organisms and detrital particles. To resolve time-lags between phytoplankton and zooplankton, we implement the life-cycle of multicellular zooplankton (here represented by copepods). The simulation successfully captures observed patterns in biomass and energy fluxes across regions. We find that carbon export correlates best with copepod biomass and trophic level, whereas the pe-ratio correlates best with the exponent of the size spectrum and sea surface temperature (SST). Community metrics performed better than NPP or SST for both deep export and pe-ratio. Time-lags between phytoplankton and copepods did not strongly affect export or pe-ratio. We conclude by discussing how can we reconcile size-spectrum theory with field sampling.

**Plain Language Summary:** Plankton are tiny but extremely abundant aquatic organisms. Plankton lock CO_2_ away from the atmosphere as they sink to the deep ocean, where carbon can be stored for hundreds of years. However, how much carbon is locked away and for how long depends on how organisms eat, defecate, and respire. We argue that these processes are reflected in the size composition of the plankton community. The size composition shows a clear relationship between the number of organisms and their body-size. The steepness of this “size-abundance relationship” describes the balance between small vs. large organisms, and has been argued to reflect how energy is transferred from small to large organisms. Since large organisms create fast-sinking particles, the size-abundance relationship could be used to estimate how much carbon is being stored in the deep ocean. Here we use a computer simulation of the global plankton community to investigate how the removal of carbon relates to the plankton community and the steepness of the size-abundance relationship. The model successfully captures patterns observed in nature. We found that the size-abundance relationship, together with the quantity of large zooplankton better explained carbon export than other measures typically used, such as photosynthesis and temperature.

**Key Points:**

- We use a global mechanistic size-spectrum model to investigate the relation between particulate export and plankton community metrics.
- We find a good correlation between export efficiency and the exponent of the size spectrum.
- Total carbon export correlated well with copepod biomass and trophic level of active copepods in the model.

## 1 Introduction

Plankton contribute to the removal of atmospheric CO_2_ by photosynthesizing in the surface ocean and sinking into the deep ocean, where remineralized carbon may remain sequestered for hundreds of years (Longhurst & Harrison, 1989; Ducklow et al., 2001). The amount of carbon exported and carbon export efficiency emerge from intricate processes that result in either carbon being respired in the surface ocean – and therefore not sequestered – or exported and respired in the deep ocean. Where and how much carbon is respired depends on the community composition and interactions between organisms who eat, respire, and excrete this carbon several times as energy flows across the foodweb. However, due to the large amount of players and processes that alter carbon export, global estimates of the flux out of the euphotic zone are highly uncertain, ranging ≈ from 3 to 12 PgC year^−1^ (Dunne et al., 2005; S. Henson et al., 2011; DeVries & Weber, 2017).

Community composition and interactions between organisms drive carbon export and its efficiency (Ducklow et al., 2001; S. Henson et al., 2019). In general, food-webs that are dominated by large organisms are expected to efficiently export large amounts of carbon (Wassmann, 1997; Stamieszkin et al., 2015). This is because large organisms produce fast-sinking particles (Small et al., 1979). These food-webs tend to be short, where NPP efficiently reaches large organisms (Wassmann, 1997). Conversely, food-webs dominated by small organisms tend to be long, with many trophic transfers. Each trophic transfer results in respiration losses, and therefore long food-webs with many trophic levels result in carbon being exported inefficiently (Wassmann, 1997).

Time-lags between phytoplankton and zooplankton are another factor that has been suggested to affect carbon export (Parsons, 1988; S. A. Henson et al., 2015; S. Henson et al., 2019). These time-lags result from the slower demographic response of multicellular zooplankton (e.g. copepods) relative to phytoplankton growth rate. Multicellular zooplankton need to grow in body size before being able to reproduce. This ontogenetic growth prevents multicellular zooplankton populations to grow as fast as phytoplankton that grow by cell division. In contrast, unicellular zooplankton (that also grow by cell division) are able to tightly follow phytoplankton dynamics. Grazing by unicellular zooplankton often results in low export efficiencies, as they contribute to long foodwebs dominated by small organisms (McNair et al., 2021), where most carbon is respired in the surface ocean. Hence, differences in life-history strategies between prey and predators can alter the amount of carbon exported.

Food-web structure, organismal size distributions, and the life cycle of organisms are therefore important factors contributing to carbon export and its efficiency. However, most models that simulate carbon export have similar simple food-web configurations. These food-web configurations tend to resolve a small and a large group of each component of the ecosystem: phytoplankton, zooplankton, and detritus (e.g. Laws et al., 2000; Siegel et al., 2014; S. A. Henson et al., 2015; DeVries & Weber, 2017; Bisson et al., 2020). These food-webs have fixed interactions, where the small/large zooplankton eats the small/large phytoplankton (and perhaps the large zooplankton also eats the small zooplankton). Yet, marine systems tend to form size-spectra with complex interactions (Sprules & Munawar, 1986; Sprules & Barth, 2016; Hartvig et al., 2011). Organisms of the same size can occupy different trophic levels, or the same organism can be at a different trophic level depending on the environmental conditions. In addition, in these models, no life cycle differences are made between zooplankton groups, preventing time-lags between prey and predators. Simple food-web configurations are convenient to understand some of the main interactions, but also miss several of the factors mentioned above. Hence, incorporating flexible food-web configurations, life histories, and size-spectra in ecosystem models might help identify new processes driving carbon export and its efficiency.

A major factor shaping marine food-webs is body-size (Hartvig et al., 2011; Andersen et al., 2016). Predator-prey interactions are size-dependent, where typically large eats small, and metabolic processes follow allometric relationships (Kiørboe & Hirst, 2014). In marine systems, the combination of these processes results in body-mass normalized size-spectra closely resembling power-law functions (*B* = *κm*^*λ*^), with varying coefficient (*κ*) and exponent (*λ*) (Sprules & Barth, 2016; Andersen, 2019). Differences in the coefficient indicate differences in the bulk biomass, whereas differences in the exponent show changes in the balance of small vs. large organisms, reflecting how efficiently energy and biomass reach larger organisms (Andersen et al., 2009). Among the emergent size spectra, low exponents (steeper spectra) indicate that energy is inefficiently channeled towards large organisms (inefficient food-webs), while communities with high exponents (flatter spectra) efficiently channel NPP to large organisms (efficient food-webs). The exponent of the size spectrum is thus a good indicator of food-web efficiency, and is therefore a potentially good indicator of carbon export and its efficiency.

Here we seek to understand how carbon export and its efficiency relate to community composition, food-web structure, size spectra, and trophic interactions between prey and predators. To do so, we use a mechanistic model of the planktonic community coupled to a 3D representation of a global ocean circulation model. We use the NutrientUnicellular-Multicellular (NUM) size-spectrum model of the planktonic community (Serra-Pompei et al., 2020). This framework is built upon the main processes at the individual level: physiology and prey size preference and encounter. The life cycle of multicellular zooplankton is also resolved, differentiating them from unicellular zooplankton. The model yields size spectra of plankton and detrital particles, which are important to determine particle sinking rates. Overall, food-web structure and the resulting particle export are emergent properties from biological interactions between organisms and the environment.

## 2 Methods

The NUM framework is a mechanistic size- and trait-based model of the planktonic community (Serra-Pompei et al., 2020). The original model resolves the size distribution of unicellular protists (autotrophic, mixotrophic, heterotrophic), the copepod community, copepod fecal pellets, and one pool of nitrogen. Here, the model has been extended to account for the size-distribution of dead cells and dead copepods, together referred to as deadfalls. The ecological model is embedded in a 3D transport matrix that represents advection and mixing of the ocean physical environment (Khatiwala, 2007). Here, we briefly explain the model and illustrate the main concepts (Fig. 1). A detailed explanation of the model and its equations can be found in the supplementary material and in (Serra-Pompei et al., 2020).

**Figure 1.**
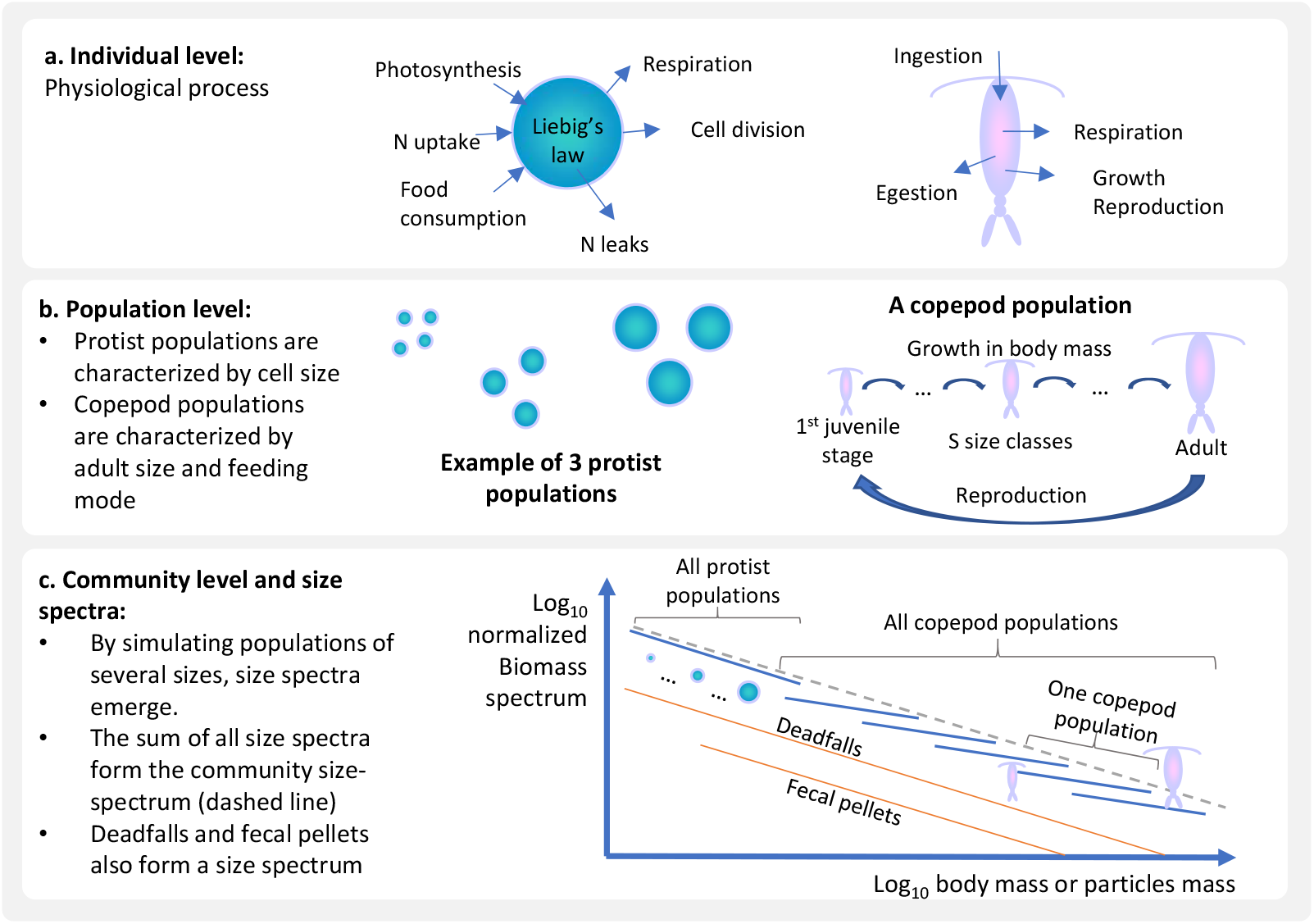
Diagram of the ecological model. (a) Community level processes are scaled from rates at the individual level, which depend on resources and prey availability as well as temperature and organism size. (b) A population is the combination of organisms that have the same trait combinations, here cell mass for protists and adult body mass and feeding mode (active vs. passive) for copepods (b). Finally, (c) the combination of all populations results in community-level processes and the emergence of size-spectra.

### 2.1 Ecological model

The model is mechanistic, where we use empirically demonstrated mechanisms at the individual level to scale to the population, community and ecosystem levels. The model generates a community of protists, copepods, fecal pellets, and deadfalls (Fig. 1c). To obtain the community size-spectrum, the model simulates several size-classes of each compartment (Fig. 1b). Protists are discretized in populations characterised by the organism’s size. Copepods also have several populations, each characterised by the adult bodymass and feeding mode. Each copepod population grows in body-size as they mature from nauplii to adults that can reproduce (Fig. 1b). The growth from nauplii to adulthood results in changes of up to two orders of magnitude in body mass. Copepods produce fecal pellets that are proportional to the organism size. Finally, both protists that die through viral lysis and copepods that die through non-consumptive mortality result in deadfalls of sizes that depend on the size of the producer. Therefore, size is the main trait describing organisms and particles, and physiological rates, predator-prey interactions, and sinking rates of particles are all size-dependent.

We consider different protist trophic strategies and copepod feeding modes. Here, protists are mixotrophic “generalists” (Fig. 1a); i.e. they can simultaneously photosynthesize, take up dissolved nutrients, and eat other organisms. Size resolves the emergence of the distinct trophic strategies across the protist size spectrum (Chakraborty et al., 2017). For example, since the smallest protists don’t have prey to eat and have a competitive advantage in nitrogen uptake, they will mainly be autotrophs. On the other hand, there is more prey available for large protists, and therefore they will tend to be heterotrophs. Intermediate sized protists will tend to be mixotrophs. Still, environmemtal conditions and prey availability will define the best trophic strategy for each size-class. As for copepods, we make a distinction between “active” and “passive” feeding modes. Active cope-pods include cruising copepods and feeding-current feeders that encompass most calanoid copepods. Passive feeding copepods are ambush “sit-and-wait” feeders that include some calanoids and most cyclopoids. Active feeding copepods constantly search for food, have high metabolic expenditures, and are more easily detected by predators. Conversely, passive feeders avoid predation by waiting for their prey to come, resulting in a lower availability of prey. These two feeding modes include the feeding strategies of most pelagic copepods.

Organisms in the model interact through competition and predation (Fig. 1). Copepods feed on protists, on other copepods, and on deadfalls and fecal pellets. Protists feed on other protists, but also have the ability to photosynthesise and take up dissolved nitrogen. Food that is not assimilated by copepods is egested as fecal pellets. The dead cells/bodies of organisms that die through viral lysis or other background mortality enter the deadfalls compartment. Deadfalls are remineralized and can be eaten by copepods. The sinking rate of fecal pellets and deadfalls is size-dependent. Overall, rather than being prescribed, the food-web configuration and resulting community trait-composition emerge from the environmental forcing (nitrogen, light, temperature), and the interactions of competition and predation.

### 2.2 Biomass spectrum

From the biomass in each size-class we obtain a size distribution of the biomass. The normalized biomass spectrum results from dividing the biomass in each size range by the size-range itself. For example, the size spectrum of protists is *P*_*k*,spec_ = *P*_*k*_*/*Δ_*P*_, and thus the unit of the biomass spectrum becomes mgC m^*−*3^ μgC^−1^ (where mgC m^*−*3^ corresponds to the biomass concentration in the water and μgC^−1^ to the bin width of the body-size range). The community size-spectrum is the sum of all the size-spectra. This normalization allows comparison between compartments, even when bin-sizes differ (see Sprules & Barth, 2016 and chapter 2 of Andersen, 2019 for more explanations regarding size-spectra conversions).

### 2.2 Ocean circulation and environmental forcing

The NUM framework is embedded within a representation of the global ocean circulation, using the “transport matrix method” (Khatiwala et al., 2005; Khatiwala, 2007). The transport matrix is derived from a coarse resolution (2.8° × 2.8°, 15 vertical levels), monthly-averaged simulation of the MITgcm (http://kelvin.earth.ox.ac.uk/spk/Research/TMM/TransportMatrixConfigs, as used in Dutkiewicz, Follows, & Parekh, 2005). The coarse resolution results in the euphotic zone being resolved in only two or three layers of the transport matrix. The temperature forcing is monthly averaged. Irradiance at the ocean surface was taken from http://sites.science.oregonstate.edu/ocean.productivity/index.php. The data was afterwards interpolated to fit the grid of the transport matrix.

### 2.4 Carbon export and carbon export efficiency (pe-ratio)

The pe-ratio is defined as the fraction of depth-integrated NPP exported as sinking particles at a given depth horizon. For both carbon export and pe-ratio, the depth horizons used in this study are 120 m and 1080 m, which are the bottom of the second and the seventh layer of the transport matrix, respectively. We consider the annual and seasonal pe-ratio. The annual pe-ratio is the ratio between NPP and export flux, both integrated over a year. The seasonal pe-ratio is the daily particle flux divided by the two weeks averaged NPP prior to export.

### 2.5 Numerics

The model configuration is flexible and any reasonable number of state variables can be implemented. Here, we use 14 protists size-classes (ranging from 10^−7^ μgC to 10^−1.5^ μgC per cell), and 8 copepod populations (6 populations of active feeding copepods and 2 populations of passive feeding copepods). Copepods range from 10^−3^ μgC for the smallest nauplii to 10^4^ μgC for the largest adult copepod. Each copepod population is divided into 5 size-classes going from nauplii to adults. There are 8 size classes of fecal pellets and 8 size-classes of deadfalls. There is one pool of dissolved nitrogen. The model was implemented in MATLAB and run for 15 years. In this run-time, the internal nutrient dynamics does not reach equilibrium. This would need much longer, unaffordable run times. To compensate for this, we initiate the nitrogen concentration with nitrate data from the World Ocean Atlas. By the end of the simulation, most model compartments reach equilibrium, yet a small drift is present due to the internal nutrient dynamics. The code can be found in the following GitHub repository https://github.com/cam-sp/NUMmodelglobal.git.

### 2.6 Model testing

Outputs of the model are compared with field data extracted from the literature. Protists biomass is compared with nanoand microplankton data from the upper 50 m of the water column from three Atlantic Meridional Transects (AMT 12-14, in summer and autumn, San Martin et al. (2006)). Copepod biomass is compared with data from the AMT 13 transect (López & Anadón, 2008). López and Anadón (2008) used a small mesh size and therefore included the smaller size range, which is often omitted in other studies. To calculate the copepod biomass we multiplied the average copepod body-mass by the abundance within each size-range of the study. Exponents of the size spectrum can also be found in San Martin et al. (2006), where they fitted a size-spectrum to nano- and micro-plankton together with mesozooplankton data. To compare net primary production (NPP) we use the data-set collected in Saba et al. (2011), where NPP field data were collected from different campaigns. We show the root mean square difference (RMSD) of NPP for each region to compare with the values obtained in Saba et al. (2011), where RMSD is:

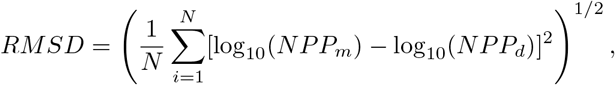

where *N* is the number of data points, *NPP*_*m*_ the modeled NPP, and *NPP*_*d*_ the data values.

For particle export, we use the data compiled in Le Moigne et al. (2013) and extended in S. Henson et al. (2019). In this data-set, the particle export is estimated between 100-200 m depth by the ^234^Th technique. Data points of export that fell within the same bin and day of the transport matrix were averaged, and the variability was shown with error bars in the figure. The same procedure was done for the NPP data. Finally, we do not compare modelled pe-ratios with the ones derived in other studies. In other studies pe-ratios are often obtained by using NPP models (from remote sensing) and differences might emerge due to model differences. We therefore stick to validate NPP and export separately without checking pe-ratios from other studies.

## 3 Results

We start by describing the general trends of biomass and energy fluxes in the foodweb while comparing them with data. Next, we investigate the drivers of food-web configurations and associated particle export and pe-ratio. All the fluxes discussed are yearly-integrated, except in the last section (3.6) where we consider seasonality.

### 3.1 Biomass

Protist and copepod biomass follow the same global trend (Fig. 2): high in temperate and sub-polar regions, and lowest in oligotrophic gyres. Compared with latitudinal field-data of nano- and microplankon biomass (Fig. 3), the model falls within observed biomass ranges in the AMT 12 transect (Fig. 3a), and in the northern hemisphere of the AMT 13 transect (Fig. 3b). However, in the latter transect, the model overestimates biomass in the southern hemisphere. This overestimation is probably due to the lack of iron limitation in the model. The model also overestimates protist biomass in the AMT 14 transect despite following well the trend (Fig. 3c). However, the biomass data from this transect is much lower than in the other two transects. Overall, despite the overestimation in some regions, modeled protist biomass follows well the general trend and magnitude of the data.

**Figure 2.**
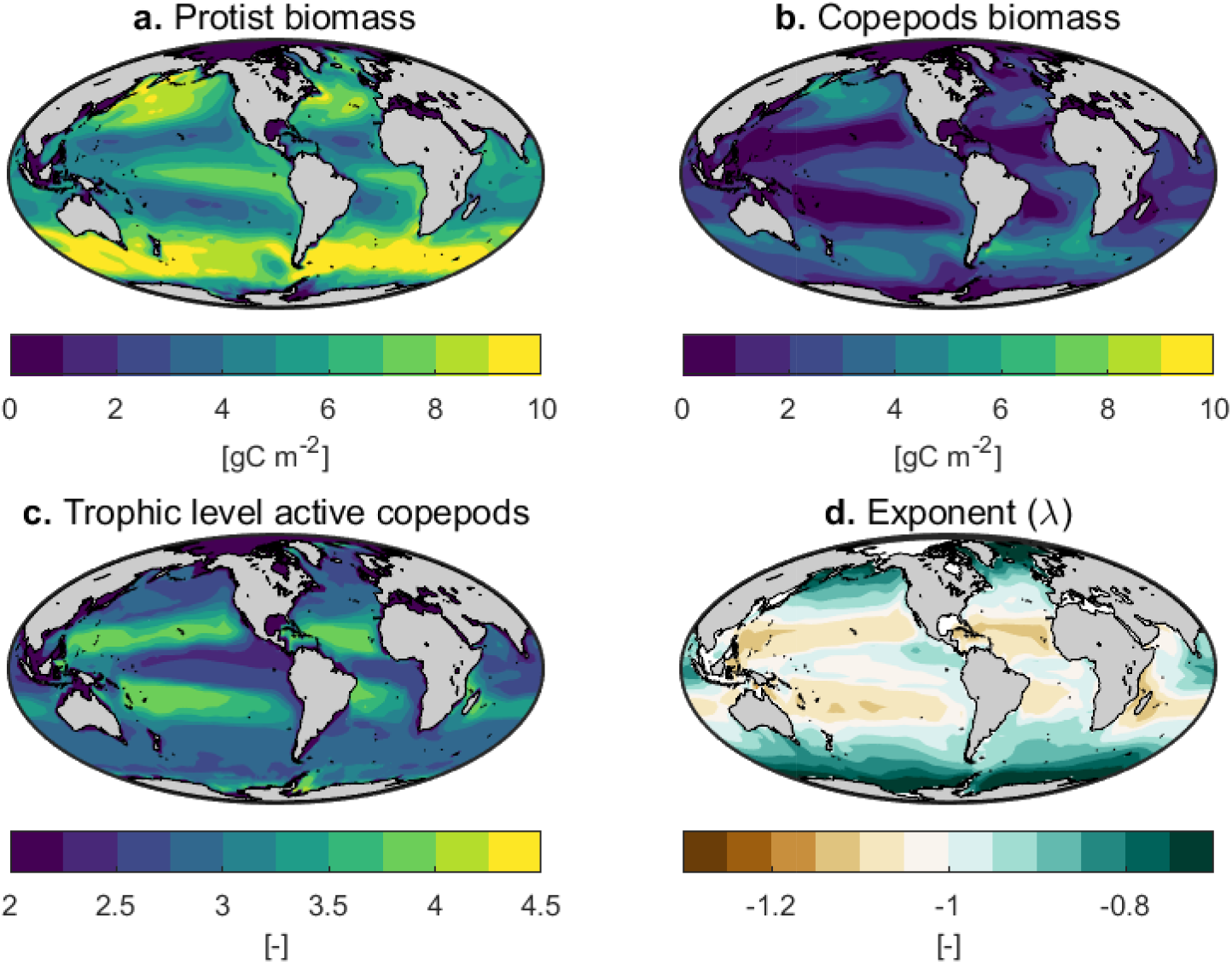
Model predictions: yearly averaged biomass of **(a)** protists and **(b)** copepods. **(c)** average trophic level of active copepods averaged over the year (trophic level is calculated as explained in the SI section B). **(d)** Yearly averaged exponent of the community size-spectrum (see figure D.1 in the SI explaining how we fit the size spectrum and obtain the parameters).

**Figure 3.**
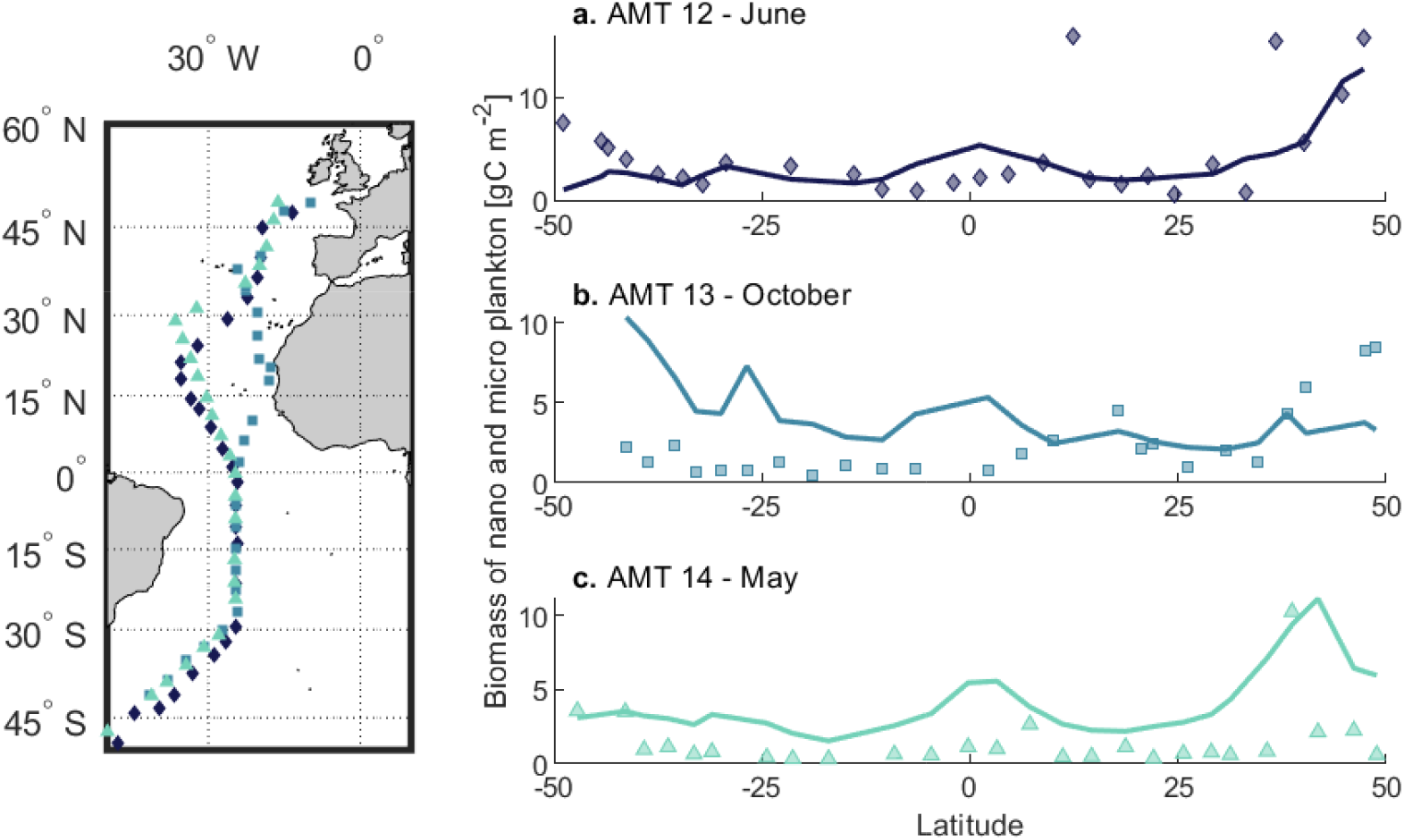
Nano- and micro-plankton biomass from the model (continuous lines) versus data (markers), both integrated over the upper 50 m. Biomass data is from three Atlantic Meridional Transects (AMT): **(a)** AMT 12 (May-June 2003), **(b)** AMT 13 (September-October 2003), and **(c)** AMT 14 (April-June 2004). Left panel shows stations where the data was collected. Data extracted from San Martin et al. (2006).

Copepod biomass trends match well the data from the AMT transect (Fig. 4), despite an overestimation between latitudes -30 and -10 (stations 22 to 18). The latter follow the protist biomass trend that was also overestimated in those regions during October. Relative copepod biomass within body-size ranges is somewhat constant across latitudes, with a dominance of small copepods in most regions (Fig. 4). The model simulates more small copepods than observed. Nevertheless, large copepods are present in most regions, including low productive regions, both in the data and the model.

**Figure 4.**
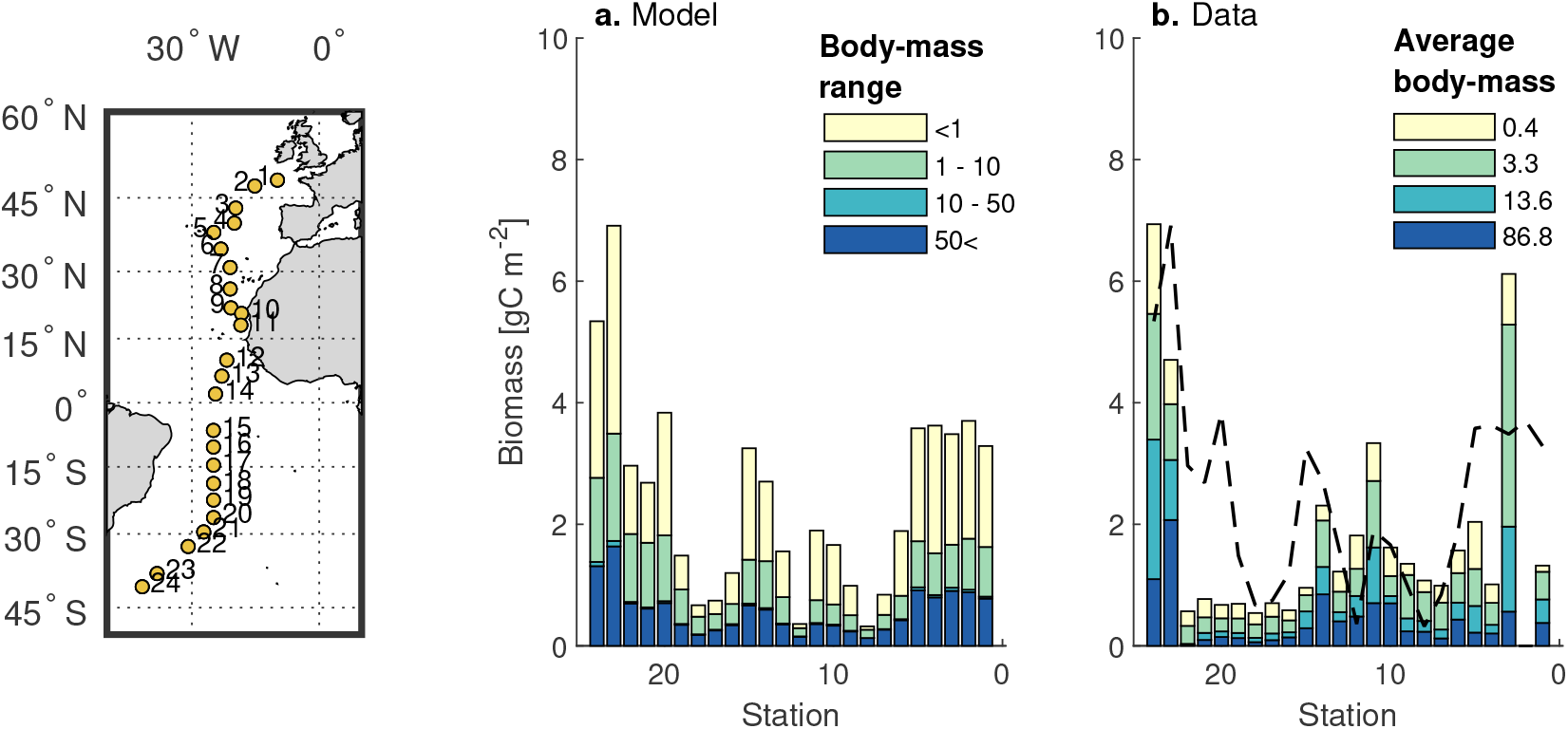
Copepod biomass along the Atlantic Meridional Transect for different body-size ranges (September-October). (a) Model estimates. (b) Data from López and Anadón (2008), obtained by multiplying the abundance by the average body-mass within each size range from field data. The smallest size-class combines adult copepods and nauplii. For easier comparison between simulated and observed data, the dashed line in panel b is the total copepod biomass from the model (from panel a). Left panel shows the stations where the data was collected in López and Anadón (2008).

### 3.2 Size spectrum

The model produces a community size spectrum (Fig. 5). We fitted a power-law function (*B*_*sp*_ = *κm*^*λ*^) to the normalized community biomass spectrum to obtain the coefficient (*κ*) and the exponent (*λ*). The yearly averaged exponent varies between *−*1.3 and *−*0.7. the exponent is always negative and always close to the theoretical value of *−*1 (Andersen & Beyer, 2006). The lowest values (steeper slopes) appear in oligotrophic regions, and the largest values in productive regions (Fig. 2). Exponents obtained from field data are also close to *−*1 (Fig. 6). Our model fits well data trends at low latitudes, but overestimates the slope at high latitudes. The data yields lower exponents at latitudes close to *−*50 and 50, particularly during the summer cruises (AMT 12 and 14, Fig. 6a,c). For the southern regions this could again be explained by iron limitation. For the northern regions, a possible reason is the presence of copepods that perform seasonal migrations, which we do not have in the model. In this case, these copepods would be present earlier in the year and deplete the system faster than in our model. In any case, both the data and model show an exponent always close to the theoretical prediction of *−*1.

**Figure 5.**
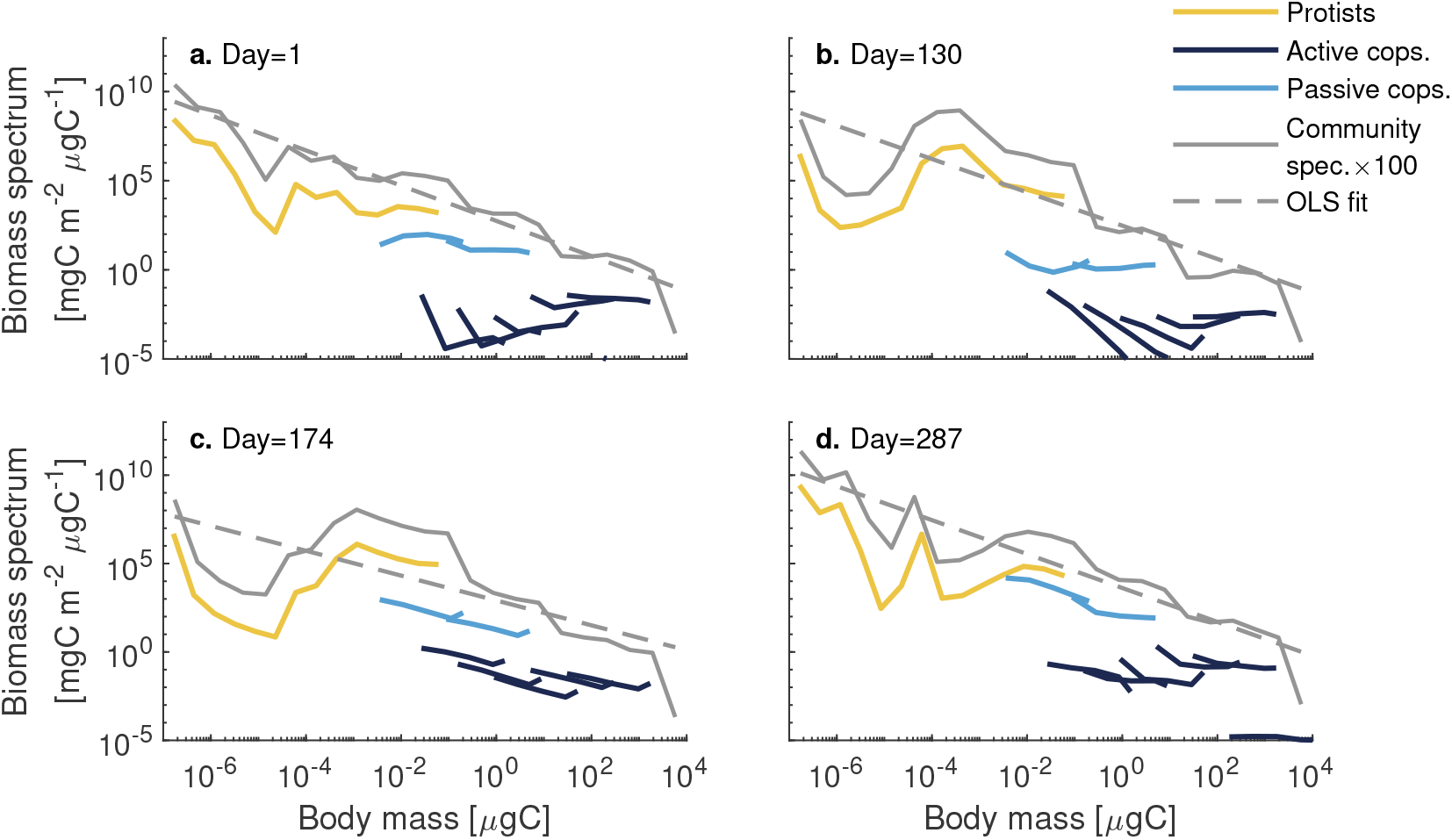
Normalized biomass spectra of the community at different times of the year in the North Atlantic: **(a)** winter, **(b)** spring, **(c)** summer, **(d)** autumn. Protists (yellow), active copepods (dark blue), passive copepods (light blue). Biomass spectra of the community (grey, we multiplied it by 100 for better visualisation), and least square fit (grey dashed line). To calculate the community spectrum, the community was divided into 24 logarithmic evenly distributed mass groups, and therefore the resolution is coarser than the spectrum within each group. The community spectrum does not include detritus.

**Figure 6.**
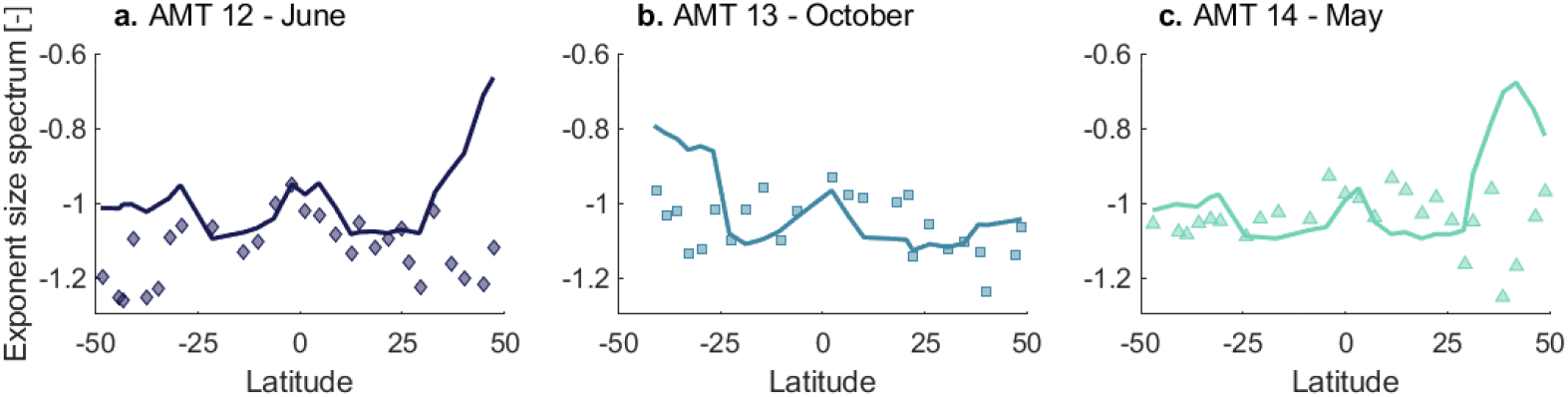
Exponent of the size spectrum from the model (continuous lines) and from field samples (markers). Data is from three Atlantic Meridional Transects (AMT): **(a)** AMT 12 (May-June 2003), **(b)** AMT 13 (September-October 2003), and **(c)** AMT 14 (April-June 2004), extracted from San Martin et al. (2006). Stations can be found in figure 3 of this paper.

Changes in the exponent seem to be driven by the presence/absence of the smallest protists size-classes and the largest copepods (Fig. 5). Some parts of the size spectrum, however, do not fit well a power-law function (e.g. Fig. 5b,c), particularly within the protists size-range. This is due to a plankton bloom, where a specific phytoplankton size group dominates the system. This highlights that a wide range of body-sizes is needed to properly fit a size-spectrum.

### 3.3 Food-web structure and trophic level

Food-web structure and trophic levels vary across regions and time (Fig. 2c and 7). The average trophic level of active copepods is highest in oligotrophic regions (Fig. 2c), where it is close to level 4, indicating long food-webs. The biomass of active copepods in these oligotrophic regions is fairly low. In temperate and sub-arctic regions, the average trophic level of active copepods decreases to between 2 and 3 (Fig. 2c and 7). This occurs when the size of primary producers increases (e.g. Fig. 7b,c versus a). Trophic levels close to 2 indicate that copepods feed mostly on primary producers, efficiently transferring energy to large organisms that produce fast-sinking particles.

**Figure 7.**
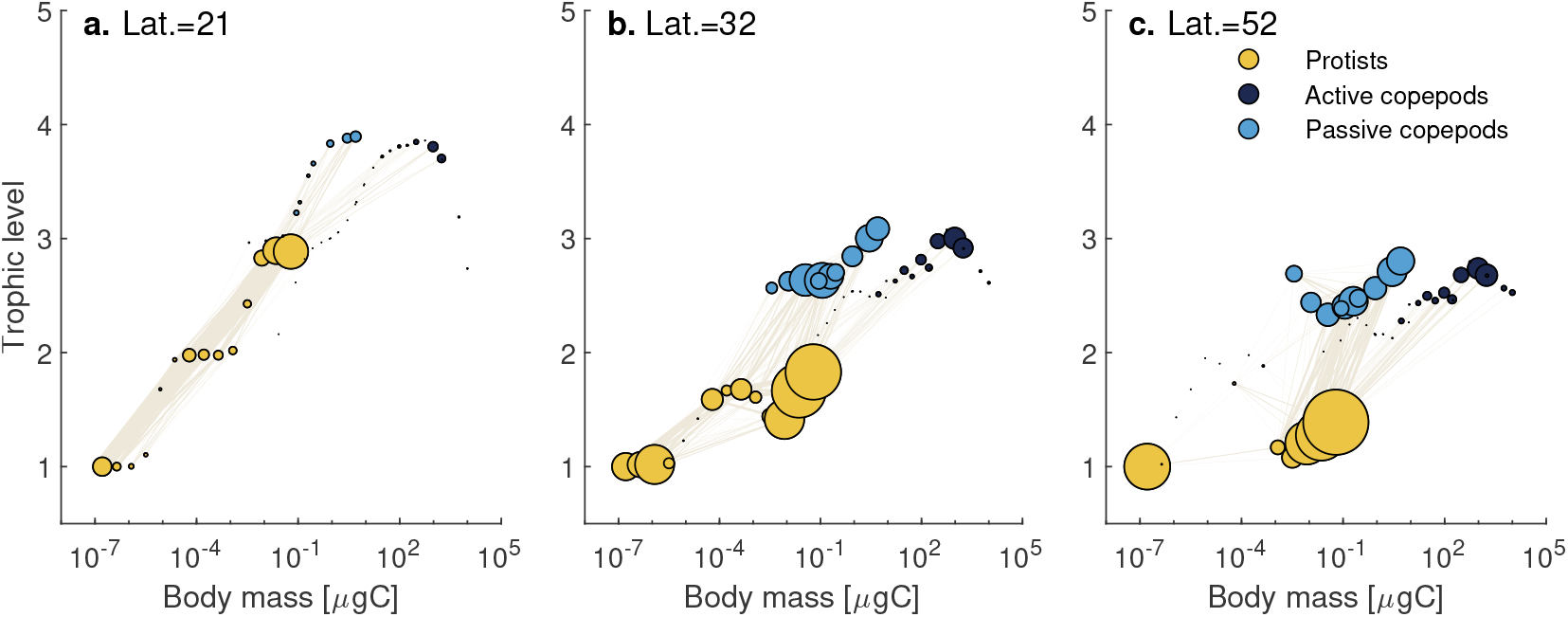
Food-webs emerging from the model shown as trophic levels (y-axis) vs. body-mass (x-axis) for three latitudes (21, 32, 52) in the North Pacific. Circle size (area) represents biomass relative to a common value for all panels. Lines connecting circles show the strength of the trophic interaction (i.e. predation, smallest values have been removed for clarity). Trophic level is calculated as explained in SI section B. Decimal trophic levels can occur due to a correction to account for mixotrophy. This also results in some large protists having a lower trophic level than their prey.

Copepods of different sizes can have similar trophic levels (Fig. 7b,c). Small passive copepods often have the same trophic level as large active copepods. This is due to the lower predator-prey mass ratio of passive feeding copepods relative to active feeders. Hence, communities dominated by passive feeders are less efficient than communities dominated by active feeders at transferring energy to large organisms and exporting carbon.

### 3.4 Primary production, carbon flux and export efficiency

Yearly averaged NPP is high in temperate and equatorial regions and lowest in the oligotrophic gyres (Fig. 8a). The annual global NPP is 62 PgC year^−1^, within the range of global NPP estimated by remote sensing (between 36 and 78 PgC year^−1^, Carr et al., 2006). Compared to field data (Fig. 9), our model performs well in some regions such as the North Atlantic and BATS but does not perform well in the Southern Ocean, where it can underestimate NPP by up to an order of magnitude. This may be due to the coarse resolution of our physical model, which results in low light levels in the surface layers. Also, NPP data in the Southern Ocean are large and variable, possibly due to some of the data being collected in a frontal zone. Still, the RMSD values in most regions are similar to or lower than the ones obtained from other NPP models (Saba et al., 2011) (note that the comparison might sometimes be imprecise due to data averaging in some bins of the transport matrix).

**Figure 8.**
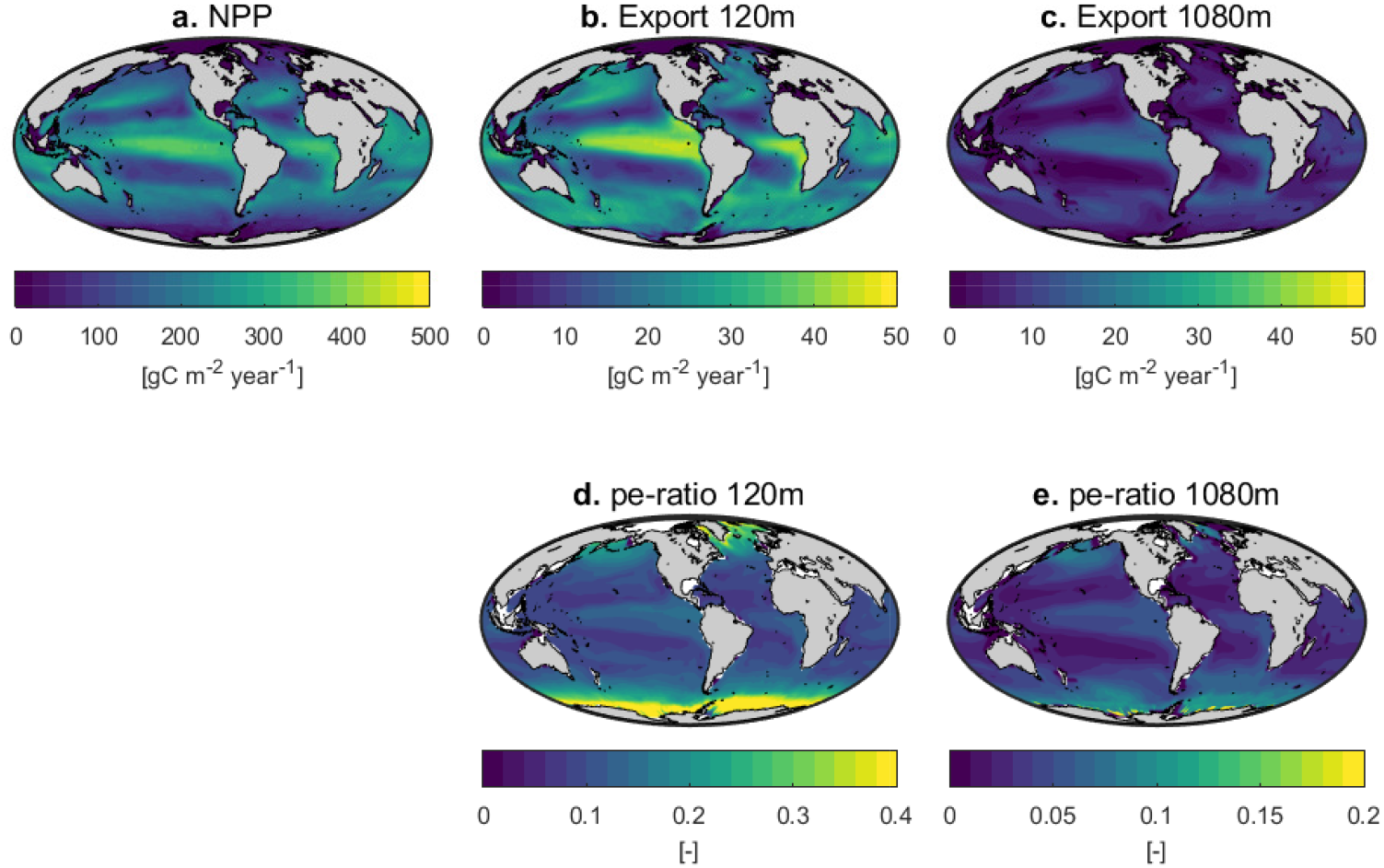
Yearly integrated **(a)** NPP, **(b)** particle export at 120 m, **(c)** particle export at 1080 m, **(d)** pe-ratio at 120 m, **(e)** pe-ratio at 1080 m.

**Figure 9.**
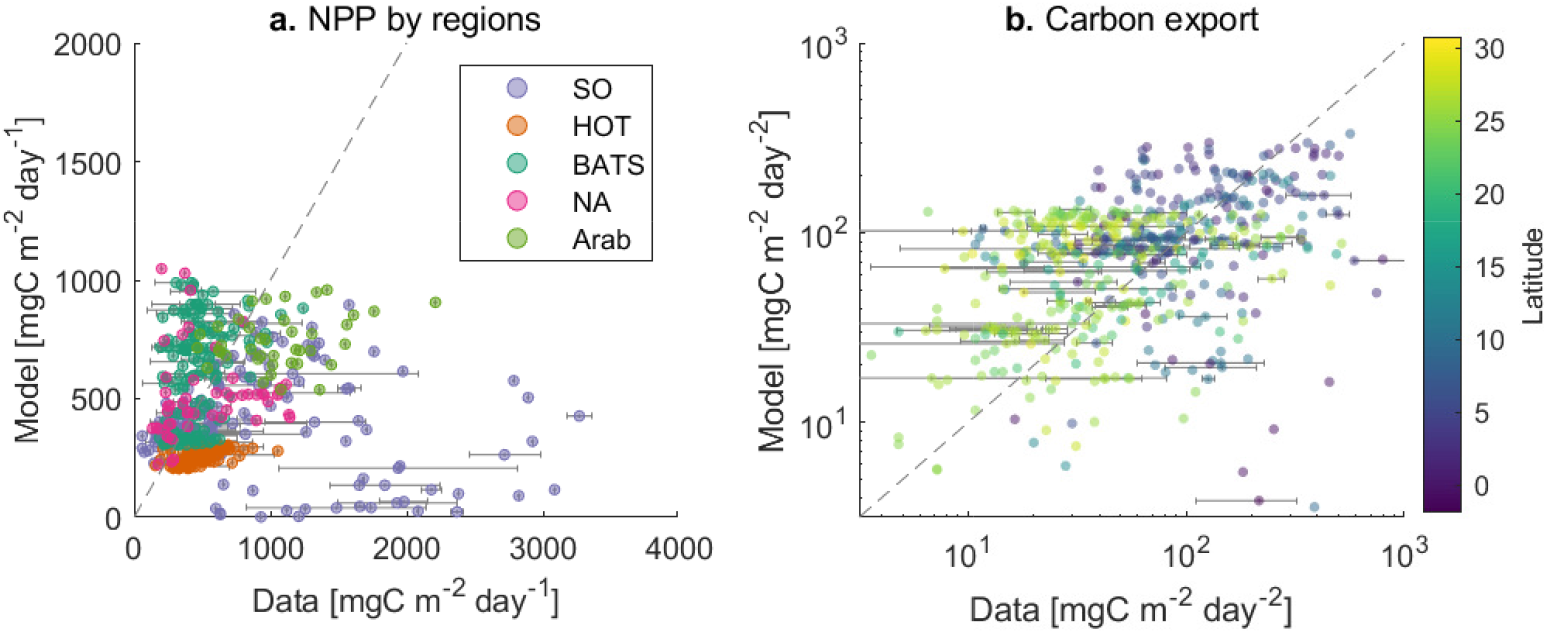
Model vs. data of **(a)** NPP and **(b)** carbon export. Colors in panel a show different ocean regions: Southern Ocean (SO, RMSD=0.53), Hawaii Ocean Time-series (HOT, RMSD=0.28), Bermuda Atlantic Time-series Study (BATS, RMSD=0.22), eastern North Atlantic (NA, RMSD=0.27), and Arabian Sea (Arab, RMSD=0.20). Colors in panel b represent latitude. Error bars appear for data that fall within the same day and bin in the transport matrix, where the shown data point is the average. NPP data was obtained from the data-set compiled in Saba et al. (2011), and carbon export data from the data-set compiled in Le Moigne et al. (2013), extended in S. Henson et al. (2019). See figures D.2 and D.3 in the Supporting Information for sampling locations.

**Figure 10.**
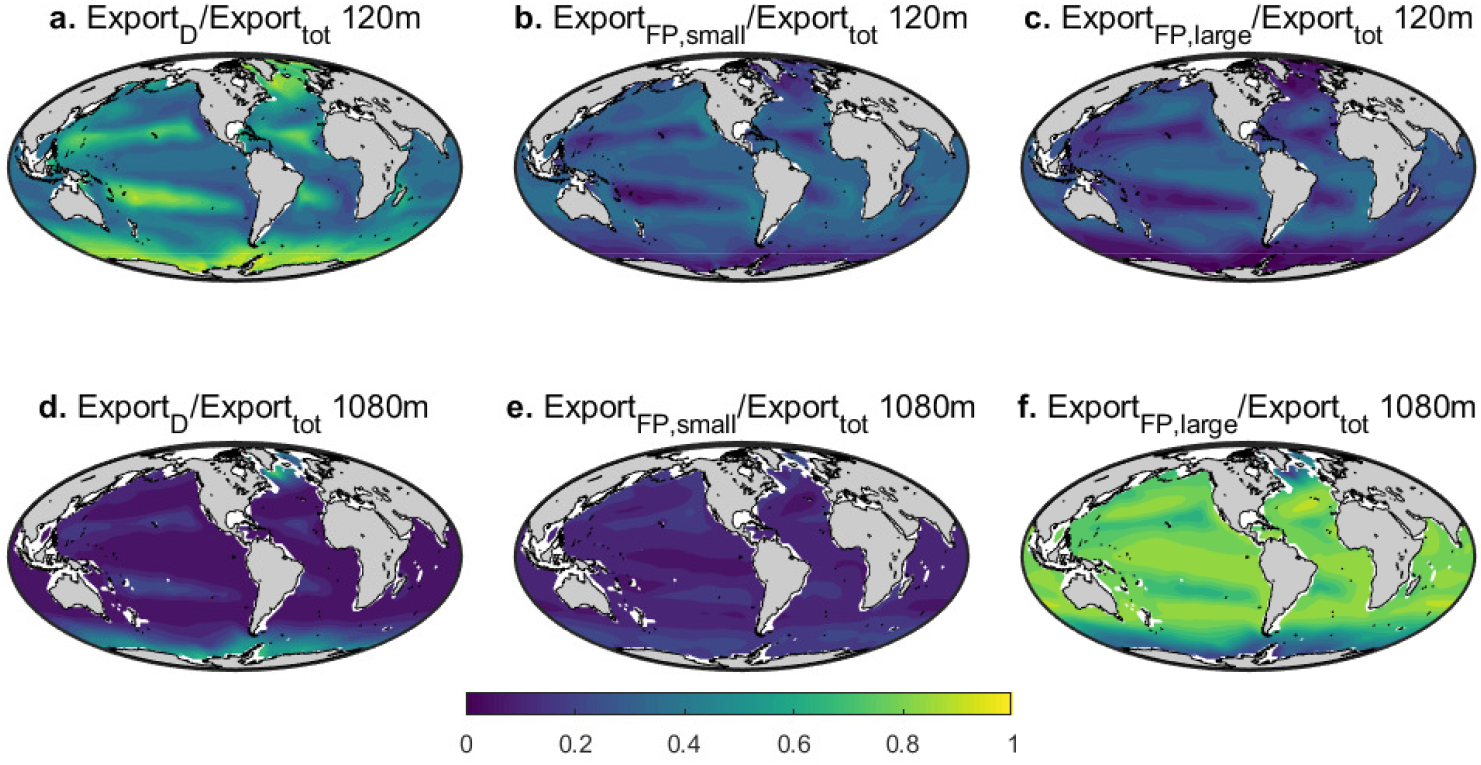
Contribution to total export from dead cells (a,d), fecal pellets from small copepods (b,e), and from large copepods(c,f) at 120 m (upper panels) and 1080 m (lower panels). Small copepods are below 1mm.

Global carbon export is 7.4 PgC year^−1^ at 120 m (and 2.0 PgC year^−1^ at 1080 m), which is within recent estimates of global carbon export (6.6 PgC year^−1^ in Siegel et al. (2014) and 9.1 PgC year^−1^ in DeVries and Weber (2017)). Yearly integrated carbon export is highest in tropical and in temperate regions (Fig. 8b and c). Compared to field data of ^234^Th measured at depths between 80 and 150 m (Fig. 9b), the spread is rather large. The yearly pe-ratio at 120 m ranges between *≈*0 in oligotrophic regions to more than 0.4 in polar regions, and pe-ratio at 1080 m is below 0.1 in most regions except in polar regions, where it reaches almost 0.2 (Fig. 8d,e).

The composition of the sinking material varies across regions and depth (Fig. 10). Deadfalls dominate export at 120 m in oligotrophic regions, in the North Atlantic, and in the Southern Ocean (Fig. 10a). Elsewhere, deadfalls and copepods fecal pellets contribute similarly to the surface carbon flux (Fig. 10b,c). However, deep export (1080 m), is dominated by fecal pellets produced by large copepods, except in the Southern Ocean and the North Atlantic. In these regions, dead cells still contribute 60% of the deep export. In conclusion, the composition of the carbon export is highly variable close to the surface, but is mainly dominated by large fecal pellets below 1000 m.

### 3.5 Relation between export, pe-ratio and community metrics

To explore the relations between community metrics and carbon export, we performed a correlation analysis of export and pe-ratio at 120 m and 1080 m with the exponent of the size spectrum, the average trophic level of active copepods, and copepod biomass. We also compared the performance of these community metrics with other more commonly used metrics such as NPP and sea surface temperature (SST, Fig. 11).

**Figure 11.**
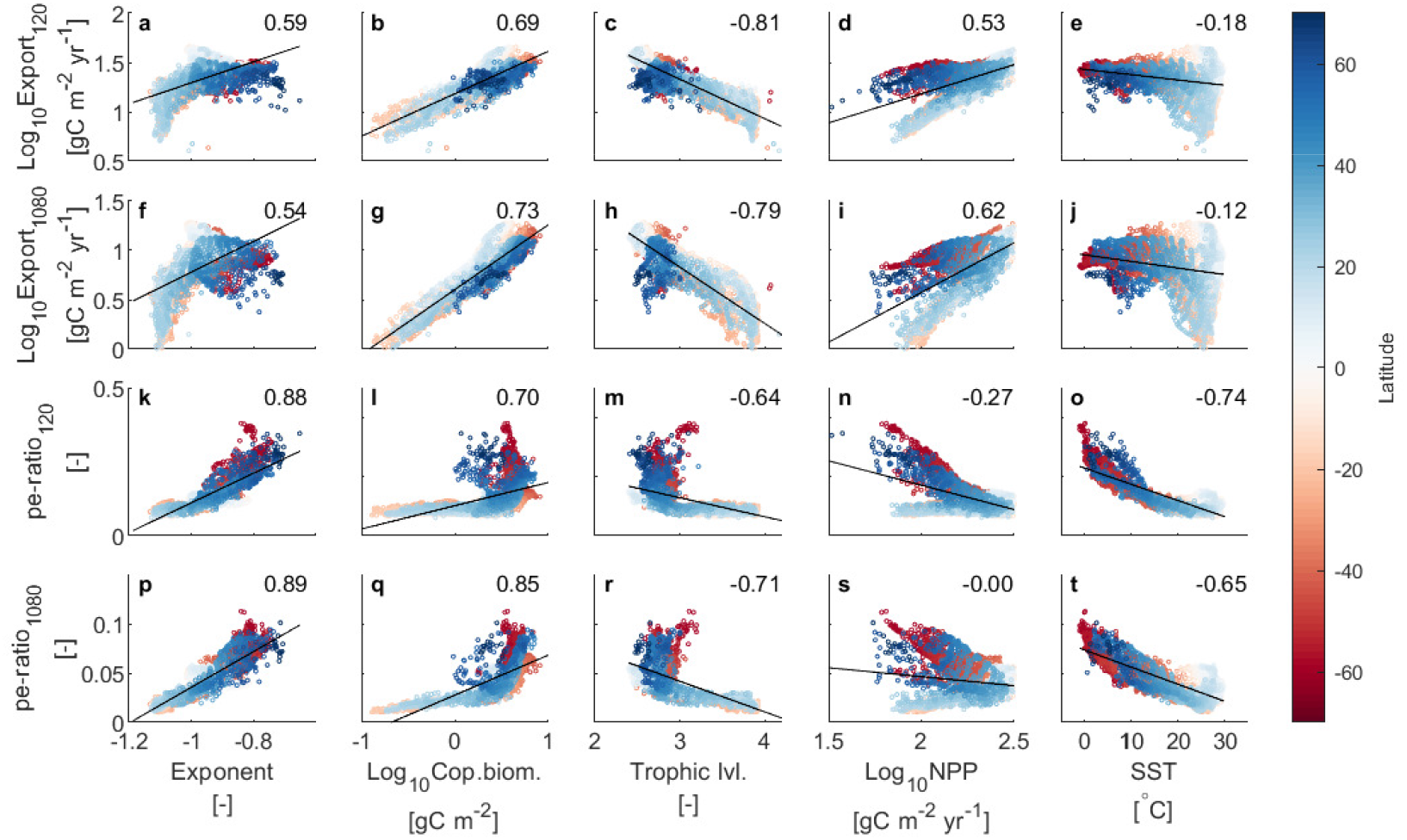
Correlation plots for export and pe-ratio at 120 m and 1080 m against the exponent of the size-spectrum (a,f,k,p), copepod biomass (b,g,l,q), average trophic level of active copepods (c,h,m,r), NPP (d,i,n,s), and SST (e,j,o,t). Rates are yearly integrated, the rest is yearly averaged. Numbers in the upper corner of each panel show the Spearman correlation coefficient.

The exponent of the size spectrum correlates strongly with pe-ratio at all depths (Fig. 11k,p), and with surface export (Fig. 11a). Large exponents (“flatter” spectra) result in high and efficient export, whereas low exponents (steeper spectra) result in low and inefficient export. That is because the exponent of the size spectrum provides information on how efficiently energy reaches large organisms that are efficient carbon exporters.

Copepod biomass correlates well with carbon export (Fig. 11b,g). This is not surprising, since in the model, copepods produce faster-sinking particles relative to protists. Due to the relative constant size-distribution of copepods, an increase in copepod biomass also incurs an increase in large copepods that contribute the most to deep export. On the other hand, surface pe-ratio is mainly unrelated to copepod biomass (Fig. 11l). In this case, there is a lower bound of copepod biomass until which pe-ratio and copepod biomass are uncorrelated, and after this threshold is surpassed, any kind of pe-ratio can be found. This effect is smoothed for the deep pe-ratio, where there is again a stronger relationship with copepod biomass (Fig. 11q).

The average trophic level of active copepods has the strongest (negative) correlation with carbon export (Fig. 11c,h). The pe-ratio shows no clear relation with trophic level (Fig. 11m,r), similar to the relationship described for copepod biomass.

NPP shows two trends with carbon export (Fig. 11d,i), one for low latitudes and another for high latitudes. On the other hand, NPP correlates negatively with pe-ratio at higher latitudes and has no clear relation at low latitudes (Fig. 11n). The variability increases for both deep export and deep pe-ratio (Fig. 11j,s), weakening the relationship between the two variables.

Finally, temperature shows no clear relation with export (Fig. 11e,j), but is strongly related to pe-ratio (Fig. 11o,t).

Overall, at the annual level, carbon export correlates best with total copepod biomass and the average trophic level of active copepods. On the other hand, pe-ratio correlates best with the exponent of the size spectrum and with temperature. Finally, NPP was strongly and negatively related to the surface pe-ratio in productive systems only. For both SST and NPP the effect weakened for deep pe-ratio.

### 3.6 Seasonality and time-lags

Until now we have considered yearly integrated rates. However, the dynamics of export and its efficiency vary over the season. We examine this in three regions with different dynamics: North Pacific, North Atlantic and an oligotrophic gyre. The North Pacific has large copepods throughout the year (Fig. 12d) and a gradual increase of phytoplankton biomass and NPP during spring (Fig. 12g). The North Atlantic has a phytoplankton bloom almost twice as intense as in the North Pacific (Fig. 12h) and copepods appear relatively late in the season (Fig. 12e). Finally, the oligotrophic gyre has a very low copepod biomass and is dominated by a microbial food-web (Fig. 12c,f).

**Figure 12.**
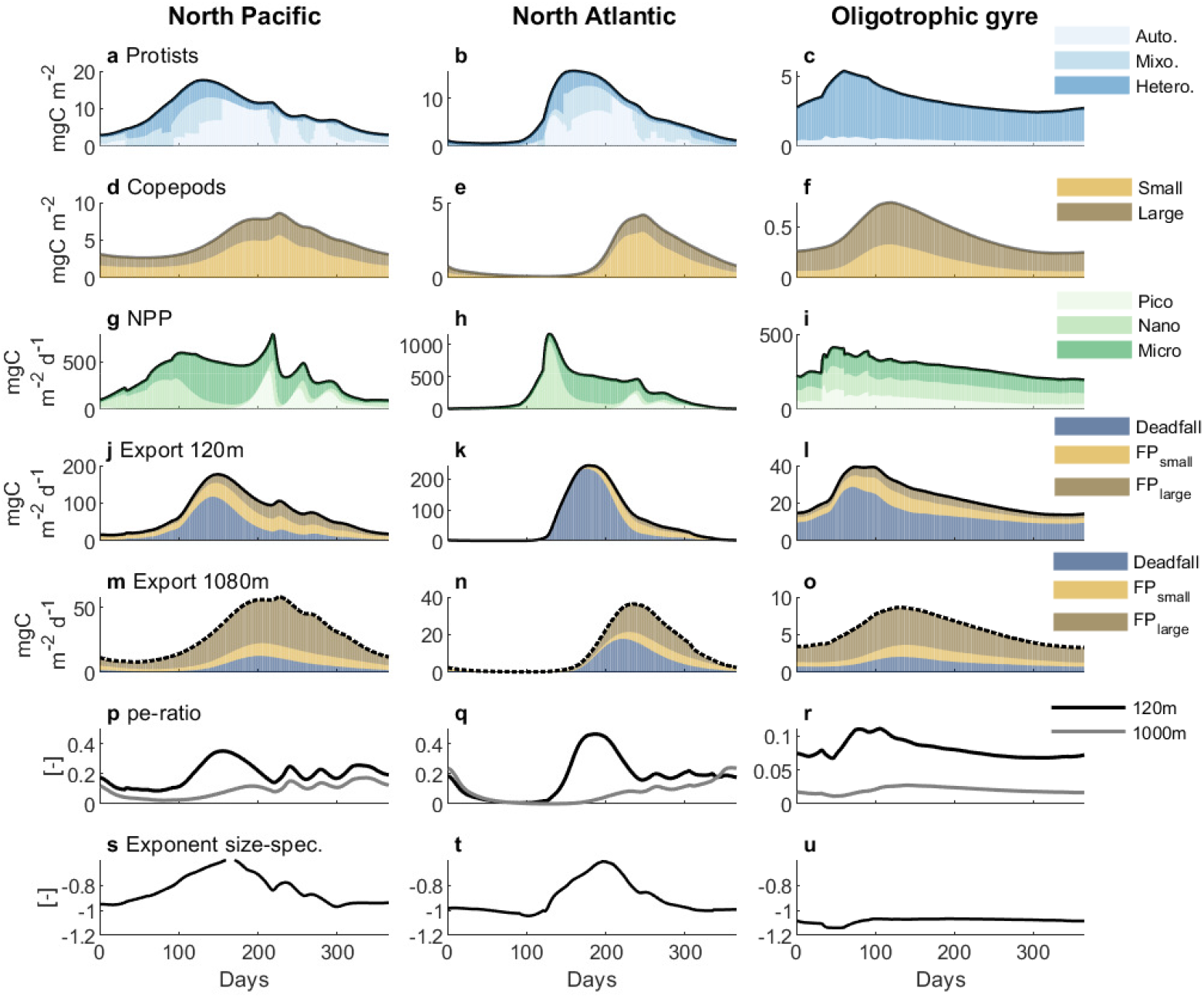
Seasonal dynamics in the North Pacific, North Atlantic and an oligotrophic region. **(a**,**b**,**c)** biomass of protists, **(d**,**e**,**f)** biomass of copepods. **(g**,**h**,**i)** NPP, **(j-o)** export at 120 m and at 1080 m, **(p-r)** pe-ratio at 120 and 1080 m and **(s-u)** exponent of the normalized size-spectrum. Note different scales on the y-axis within columns.

The different dynamics between the North Atlantic and North Pacific emerge from the differences in winter phytoplankton biomass due to deep mixing. In the North Pacific, copepods are sustained by protists biomass during the winter, and are able to control the phytoplankton spring bloom, which is therefore not very pronounced. There is still a time-lag between copepods and protists, but shorter compared to the one in the North Atlantic. The constant presence of large copepods results in carbon export being dominated by fecal pellets, even during the spring bloom. The pe-ratio is high and follows the dynamics of export, particularly just after the bloom when NPP decreases while export is still high.

In the north Atlantic, copepod biomass is lower because copepods need to recover from the deep winter mixing. The delay between the spring bloom and peak biomass is also longer, and therefore most export is dominated by dead protists in spring during and after the bloom (Fig. 12k). In this case, the pe-ratio at 120 m is very high just after the bloom, but is heavily attenuated due to the slower sinking rates and therefore does not result in a high pe-ratio at 1000 m. Hence, time-lags affect the composition of the export flux, but not necessarily the pe-ratio. Ultimately the pe-ratio is defined by total export, the fast dynamics of NPP relative to export (e.g. after the bloom), and community composition, the latter defining the sinking rates of particles.

The exponent of the size spectrum follows surface pe-ratio trends, but does not necessarily follow the trend of deep pe-ratio (Fig. 12p-u). It thus seems that pe-ratio correlates well with variation of the exponent across regions but not necessarily over time within a specific region.

## 4 Discussion

We sought to understand how carbon export and its efficiency relate to the size spectrum and community composition of the planktonic community. These metrics included the exponent of the size spectrum, trophic level of organisms, and copepod biomass. We also wanted to understand how time-lags between primary producers and copepods affected export and its efficiency. The analysis was made with a mechanistic trait-based model that resolves the size structure of both the planktonic community and the sinking detritus. Food web structures and size spectra emerge from the model rather than being prescribed. Simulated biomass of copepods and protists follow well-observed trends from field data. The emergent food-webs and size-spectra result in differences in carbon export and its efficiency.

The main results are: (i) carbon export correlates best with copepod biomass and the average trophic level of active copepods, whereas (ii) pe-ratio correlates best with the exponent of the size spectrum and SST. (iii) These community metrics correlate better with deep export and deep pe-ratio than commonly used metrics, such as NPP or SST. Finally, (iv) time-lags between phytoplankton and copepods change the composition of the material exported, but do not strongly a_ect export or pe-ratio.

### 4.1 Can we use size spectra to estimate carbon export and its efficiency?

The modelled carbon export correlates better with the exponent of the size spectrum than with NPP or temperature. An interesting avenue could therefore be to use this metric to estimate carbon export. However, measuring a size-spectrum in the field is not easy, as it requires sampling organisms/particles that range several orders of magnitude in mass. Therefore, many studies focus on sampling only one part of the size spectrum (e.g. phytoplankton). Dynamics of specific size-groups of unicellular organisms can quickly vary in time, for example, during a phytoplankton bloom. In such conditions, a power-law will not fit the spectrum when only the phytoplankton size-range is considered (Fig. 5b,c). The irregularities observed in size-spectra (often referred as “domes”) are common (Sheldon & Parsons, 1967). These domes reflect other properties of the foodweb, such as changing top- vs. bottom-up control within a size range (Rossberg et al., 2019). Thus, size-spectrum theory applies only when fitted across a wide size-range of organisms.

Field-sampling over a large size range is possible (Lombard et al., 2019), but the space and time resolution of these measurements is scarce and the collection demanding. An approach that would overcome this limitation is to get proxies of the size-spectrum via remote sensing (Kostadinov et al., 2009), however this approach still needs to be validated. If sampling plankton size-spectra in the field becomes easier to achieve, or if good proxies of the size-spectrum are developed, size-spectra may become a powerful tool to quantify ecosystem processes such as trophic transfer efficiency to large organisms (such as fish) and carbon export to the deep ocean.

### 4.2 Export, pe-ratio, and time-lags

Time-lags between primary production and peak copepod biomass do not affect carbon export or its efficiency *per-se*. They mainly affect the composition of the material being exported. Some studies suggest that strong trophic coupling between phytoplankton and their predators can reduce export and its efficiency due to trophic transfer losses and higher remineralization rates in the surface ocean (Parsons, 1988; S. Henson et al., 2019). These studies often assume the predator to be mesozooplankton. However, here, we show that deep export is maximal when copepod biomass is high. Conversely, dominance by protists during spring blooms results in large surface export, but not deep export. We expect this latter result to change if formation of detrital aggregates was modelled. As aggregates become larger, their sinking rates increase. This may particularly happen during diatom blooms, potentially resulting in a high export and pe-ratio in “uncoupled systems”. Overall, whether export and its efficiency are high or low in coupled or uncoupled systems depend on how efficient prey are at exporting carbon relative to their predators.

Another result from the model is that surface export often lags NPP, and deep export does not follow NPP dynamics. These differences in timing between export and NPP complicates the interpretation of pe-ratio values. For instance, the pe-ratio at surface follows carbon export, whereas the deep pe-ratio is not necessarily higher when deep export is high. Rather, the deep pe-ratio increases when NPP decreases. These differences in rate of change of export end NPP have been shown to give different pe-ratios depending on how NPP is averaged (Laws & Maiti, 2019). Total export has a more intuitive dynamic than pe-ratio, since pe-ratio is a function of two rates that vary at different time-scales and where the uncertainties of both measurements are propagated. Thus, estimating total export may be more useful than attempting to estimate the pe-ratio.

### 4.3 Contribution by copepods to carbon export

Deep export in most regions is dominated by fecal pellets of large copepods, even in oligotrophic regions. This result agrees with other studies that found copepod size to be an important driver of carbon export (Stamieszkin et al., 2015). This is, however, not necessarily supported by other studies. Using field data, a recent study found regimes of low carbon export in regions where macrozooplankton biomass and bacteria were high (S. Henson et al., 2019). Other studies argue that copepods can strongly attenuate the carbon flux by fragmenting detrital particles (Wexels Riser et al., 2007, 2010; Cavan et al., 2017; Mayor et al., 2020). We consider consumption of detritus by copepods, but not a reduction in particle size due to particle fragmentation. Therefore, in the model, losses by simple trophic transfer are not enough to attenuate the flux.

### 4.4 Comparison with other models and model limitations

Our model fits biomass data well, but less so in terms of NPP and carbon export. Other models which are optimised with observations perform better at estimating these rates (Stock et al., 2014; Siegel et al., 2014; DeVries & Weber, 2017), but provide less mechanistic detail. Still, in terms of NPP, the variation found for each region is lower or similar to the values found in 21 NPP models (Saba et al., 2011), suggesting that the model performs relatively well. Similarly, our global estimate of carbon export at 100 m, 7.4 PgC year^−1^, falls within the range of estimated values of some of the most recent studies (6.6 PgC year^−1^ in Siegel et al. (2014) and 9.1 PgC year^−1^ in DeVries and Weber (2017)). It should also be kept in mind that the data used to validate models have high uncertainties. We used a global data-set where carbon export by sinking particles was measured via the ^234^Th method. However, due to adsorption of ^234^Th on filters and preferential collection of suspended versus sinking particles, this method can underestimate the carbon flux by approximately 2-fold in some regions (Quay, 1997; Buesseler et al., 2000). Moreover, the variability of export in each region is high, and often the standard deviations in each region are close or larger than the mean value (Le Moigne et al., 2013). Hence, given the uncertainty in NPP and export observations, it is hard to reliably validate model performance.

Despite the high ecological complexity of our model, the biogeochemistry is simplistic. We use nitrogen as the sole nutrient in the system, whereas most global models consider other limiting nutrients such as iron, phosphorus or silica (S. Henson et al., 2011; DeVries & Weber, 2017; Ward et al., 2018), which can be important limiting factors in some ocean regions. Therefore the model does not resolve iron limited regions (e.g. the southern ocean). In addition, the coarse resolution of the transport matrix prevented us from obtaining carbon export just below the photic layer, which has been recommended in recent studies (Buesseler et al., 2020). Instead we measured it at fixed depths (120 m and 1080 m) probably causing some biases when comparing carbon export across regions.

Protists in the model are not separated into functional groups. Instead, the trophic strategy (autotrophy, mixotrophy or heterotrophy) emerges as a function of cell size and the environment. This configuration still captures the main dynamics observed in nature: a dominance of small primary producers in oligotrophic regions, larger primary producers in more productive regions, and the constant presence of unicellular zooplankton and the microbial food-web. This simplification becomes an advantage as it captures complex dynamics while being based on a relatively low set of parameters and processes.

Other organisms that may contribute significantly to carbon export but are not included in our model are gelatinous zooplankton (Luo et al., 2020). The inclusion of these organisms would probably increase export due to (i) their large bodies that can quickly sink to the bottom, and (ii) their large predator-prey mass ratio. Large predator-prey mass ratios generate “shortcuts” in the food-web, where energy from very small organisms is efficiently transferred to larger ones, further enhancing carbon export and its efficiency. Some recent global models now include gelatinous zooplankton (Heneghan et al., 2020), but these organisms still lack in most (all?) global biogeochemical models.

Finally, we do not represent diel and seasonal vertical migrations of zooplankton. Vertical migration play an important role in the survival and life cycle of copepods, and in carbon export (Jónasdóttir et al., 2015; Hansen & Visser, 2016; Steinberg & Landry, 2017; Pinti et al., 2021). Implementing vertical migrations may be done through optimisation (e.g. Brun et al., 2019; Pinti et al., 2019; Pinti & Visser, 2019). However, implementing behaviour together with population dynamics is challenging, especially if considered at the global scale. Implementing the active export pathway is urgently needed, since other modelling studies have suggested that the active carbon flux may have been responsible for increasing the efficiency of the biological carbon pump (Fakhraee et al., 2020).

## 5 Conclusion

We have investigated how carbon export and its efficiency relate to the size spectrum and community composition of the planktonic community. We have shown that carbon export correlates well with copepod biomass and trophic level, and that pe-ratio correlates best with the exponent of the size spectrum and temperature. Community metrics correlate better with deep export and deep pe-ratio than SST and NPP. Time-lags between phytoplankton and zooplankton do not necessarily affect carbon export or its efficiency. Our framework captures complex community dynamics scaled from simple individual-level processes. The model also successfully captures observed inter-biome differences in plankton biomass and rates. This study has shown the potential of more complex ecological models to explore and understand ecosystem functions and biogeochemical processes at the global scale.

## Acknowledgments

This work was supported by the Gordon and Betty Moore Foundation through award 5479, and by the Centre for Ocean Life, a VKR Centre for Excellence funded by the Villum Foundation. CSP was also funded by the Simons Foundation Postdoctoral Fellowship in Marine Microbial Ecology. B.A.W. was funded by a Royal Society University Research Fellowship. Data-sets for this research are included in these papers (and their supplementary information files): San Martin et al. (2006); López and Anadón (2008); Saba et al. (2011); Le Moigne et al. (2013); S. Henson et al. (2019).

*from main text and SI combined

