## Supplemental information for "Linking plankton size spectra and community composition to carbon export and its efficiency"

**B Trophic level calculation** **15**

**C Net primary production** **15**

**D Supplementary figures** **16**

### A Model equations

The following sections includes an explanation of the model where the equations are compiled in tables. For a more detailed description of the model refer to Serra-Pompei et al. (2020), and consider the modifications listed below.

#### A.1 Additions and modifications to the original model

- Respiration in copepods is now dependent on the feeding level (Eq. A.3.2).
- We have added the deadfalls compartment.
- Here, the nitrogen inputs and mixing are taken care by the transport matrix, i.e. the semi-chemostat equation that was present in Serra-Pompei et al. (2020), representing nitrogen inputs and the mixed layer dynamics, is not needed here.

#### A.2 Ecological model

The compartments in the model are: copepods ($C$), protists ($P$, which can be autotrophs, mixotrophs, or heterotrophs), nitrogen ($N$), copepod fecal pellets ($F$), and dead cells by protists and copepod carcasses, which we refer as deadfalls ($D$). All compartments have several size-classes (except for the $N$ compartment). Each size-class within each compartment is simulated by an ordinary differential equation. Hence, the whole system of equations goes as follows: for each compartment ($C, P, N, F, or D$), the dynamics within each size-class $i$ are as shown in table A.1. The equations of each compartment are explained in the following sections.

| | |
|---|---|
| Copepods | $$\frac{\mathrm{d}C_{i,s}}{\mathrm{d}t} = \begin{cases} \varepsilon_{\mathrm{r}} g_S C_S + g_1 C_{i,1} - \gamma_1 C_{i,1} - \mu_1 C_1, & \text{for } s = 1 \\ \gamma_{s-1} C_{i,s-1} + g_s C_{i,s} - \gamma_s C_{i,s} - \mu_s C_{i,s}, & \text{for } 2 \leq s < S \\ \gamma_{S-1} C_{i,S-1} - \mu_S C_{i,S}, & \text{for } s = S \end{cases} \quad \text{(A.1)}$$ |
| Protists | $$\frac{\mathrm{d}P_k}{\mathrm{d}t} = \nu_{\mathrm{u},k} P_k - \mu_{\mathrm{u},k} P_k \quad \text{(A.2)}$$ |
| Fecal pellets | $$\frac{\mathrm{d}F_l}{\mathrm{d}t} = \sum_{i=1}^{I} \sum_{s=1}^{S} f_{\mathrm{pp},i,s} C_{i,s} - r F_l - \mu_{\mathrm{p,f},l} F_l \quad \text{(A.3)}$$ |
| Deadfalls (dead cells and copepods) | $$\frac{\mathrm{d}D_j}{\mathrm{d}t} = \sum_{k=1}^{K} \mu_{\mathrm{uk}} P_k + \sum_{i=1}^{I} \sum_{s=1}^{S} (\delta \mu_{\mathrm{htl}i,s} C_{i,s}) - r D_j - \mu_{\mathrm{p,D},j} D_j \quad \text{(A.4)}$$ |
| Nitrogen | $$\frac{\mathrm{d}N}{\mathrm{d}t} = \frac{1}{Q_{\mathrm{C:N}}} \Big[ r(\sum_{l=1}^{L} F_l + \sum_{j=1}^{J} D_j) + \sum_{k=1}^{K} (\eta_{\mathrm{leaks}.k} - \eta_{\mathrm{N}.k}) P_k + \eta_{\mathrm{DON},s} + \sum_{i=1}^{I} \sum_{s=1}^{S} (1-\delta) \mu_{\mathrm{htl},i,s} C_{i,s} \Big] \quad \text{(A.5)}$$ |

**Table A.1.** Equations of all the model compartments. Terms appearing in this table are specified in table A.2. Indexes are: $s$ for a copepod size-class within a population $i$, and $k, l, j$ are each size class of protists, fecal pellets and deadfalls respectively. Capital letters of the same indices represent the total number of size-classes in each compartment. For the fecal pellets and deadfalls, the sums of all the rates originating from copepods and protists are summed only for the size ranges corresponding to each size group of the detritus.

| Symbol | Description | Equation # in the SI |
|---|---|---|
| $g_s$ | Biomass accumulation within each copepod stage | Eq. A.3.4 |
| $\gamma_s$ | Maturation rate copepods | Eq. A.3.5 |
| $\mu_s$ | Total mortality rate copepods | Eq. A.3.8 |
| $\nu_{\mathrm{u}}$ | Growth rate protists | Eq. A.3.3 |
| $\mu_{\mathrm{u}}$ | Total mortality rate protists | Eq. A.5.3 |
| $f_{\mathrm{pp}}$ | Fecal pellet production by copepods | Eq. A.3.6 |
| $\mu_{\mathrm{p,f}}$ | Consumption of fecal pellets by copepods | Eq. A.7.1 |
| $r$ | Remineralization rate of fecal pellets or deadfalls | Tab. A.6 |
| $\mu_{\mathrm{htl}}$ | Mortality by higher trophic levels on copepods | Eq. A.3.11 |
| $\mu_{\mathrm{p,D}}$ | Consumption of deadfalls by copepods | Eq. A.7.2 |
| $\eta_{\mathrm{leaks}}$ | Nitrogen leaks by protists | Eq. A.5.4 |
| $\eta_{\mathrm{N}}$ | Nitrogen uptake by protists | Eq. A.5.1 |
| $\eta_{\mathrm{DON}}$ | Nitrogen excretion by copepods | Eq. A.3.7 |
| $Q_{\mathrm{C:N}}$ | Carbon to nitrogen ratio | Tab. A.6 |

**Table A.2.** Terms appearing in table A.1. All rates are in units of day$^{-1}$ except $\eta_{\mathrm{DON}}$ which is in µgC L$^{-1}$ d$^{-1}$. Equations of these rates are listed below and follow the equation numbers of the last column in this table.

#### A.3 Copepod population model

The copepod population model is a discretized version of the McKendrick–von-Foerster (MvF) equation (De Roos et al., 2008), also used in (Serra-Pompei et al., 2020). The original MvF solves in a partial differential equation the growth in body-size of a cohort and its further reproduction. To discretize it, we make approximations to solve it as a system of ordinary differential equations (De Roos et al., 2008). Here corresponding to equation A.1. The model considers individual-level processes such as reproduction, growth in body size, feeding and mortality. Growth and reproduction are food- and temperature- dependent. Copepods have determinate growth, meaning that they stop growing once maturation is reached. Hence, all the energy gained by juveniles goes into growth of body size, whereas adults invest all their energy into reproduction. Note that there is a distinction by how a population and an individual are treated: an individual is characterised by its size and feeding mode, and a population is characterised by the *adult size* and the feeding mode (i.e. within each population several individuals of all stages have different size-dependent individual rates). In the next sections, we first explain individual-level processes and next how they scale to the population level. Finally we explain the traits characterising copepods.

##### A.3.1 Individual-level processes

Equations of the copepod model can be found in table A.3. All individuals are characterised by their size $m$, they are born with size $m_b$ and mature at size $m_m$. Copepod's energy is obtained through ingestion of prey. Individuals have a size-dependent preference for prey (Eq.A.4.1). The food available for a copepod is the product of the preference function with the biomass of prey, where copepod's prey are protists, other copepods, detritus and fecal pellets (Eq. A.4.2). Prey ingestion is modelled by a Holling type II functional response that saturates at the maximum ingestion rate $h(m)$ (Eq. A.3.1). Carbon-specific growth rate is the energy available after food assimilation and respiration (Eq. A.3.3). Respiration rate (Eq. A.3.2) is a fraction of the maximum ingestion rate A.3.2, where there is a constant fraction that si assumed to be basal metabolism $k_{\mathrm{b}}$, and another fraction $k_{\mathrm{SDA}}$ that varies with the feeding level. Finally, if the net energy gain $\nu$ is positive, this energy is invested into somatic growth or reproduction (Eq. A.3.4). On the other hand, if the biomass production rate $\nu$ is negative (i.e., if respiration exceeds food assimilation), the individual suffers starvation mortality (Eq. A.3.9).

##### A.3.2 Population level

The net energy gained by individuals goes into somatic growth or reproduction. Somatic growth is modelled by a maturation function $\gamma$ (Eq. A.3.5). All juvenile stages invest their energy into somatic growth. Adults invest all their net energy into reproduction (determinate growth). The growth through stages and reproduction are modelled through a system of coupled ordinary differential

| | Units | Equation | |
|---|---|---|---|
| **Energy gain** | | | |
| Feeding level | - | $f(m) = \frac{v(m)E(m)}{v(m)E(m)+h(m)}$ | (A.3.1) |
| Respiration | $d^{-1}$ | $\kappa(m) = \epsilon h m^n (k_{\mathrm{b}} + k_{\mathrm{SDA}} f(m))$ | (A.3.2) |
| Net energy gain | $d^{-1}$ | $\nu(m) = \epsilon h m^n f(m) - \kappa(m)$ | (A.3.3) |
| Growth rate | $d^{-1}$ | $g(m) = \max(0, \nu(m))$ | (A.3.4) |
| Maturation rate | $d^{-1}$ | $\gamma_s = \frac{g_s - \mu_s}{1 - (\frac{m^+_{s-1}}{m^+_s})^{1-\mu_s/g_s}}$ | (A.3.5) |
| Fecal pellets production | $d^{-1}$ | $f_{\mathrm{pp}}(m) = (1-\epsilon) h m^n f(m)$ | (A.3.6) |
| DON leaks by copepods | µgC $L^{-1}$ $d^{-1}$ | $\eta_{DON} = \sum_i^I \sum_s^S \kappa m^p C_{i,s} + \sum_i^I (1-\varepsilon_{\mathrm{r}}) g(m_a) C_S$ | (A.3.7) |
| **Mortalities** | | | |
| Total mortality | $d^{-1}$ | $\mu(m) = \mu_{\mathrm{st}}(m) + \mu_{\mathrm{pr}}(m)$ | (A.3.8) |
| Starvation mortality | $d^{-1}$ | $\mu_{\mathrm{st}}(m) = \min(0, \nu(m))$ | (A.3.9) |
| Predation mortality | $d^{-1}$ | $\mu_{\mathrm{pr}}(m_{\mathrm{py}}) = c_{\mathrm{py}}(\tau_{\mathrm{py}}, m_{\mathrm{py}}) \sum_i^I \sum_s^S \frac{\Phi(m_{\mathrm{py}}, m_{i,s})}{E_{i,s}} h(m) f_{\omega,i}(m_{i,s}) C_{i,s}$ | (A.3.10) |
| Mortality by higher trophic levels | $d^{-1}$ | $\mu_{\mathrm{htl}}(m) = p_{\mathrm{htl}}(m) \frac{\mu_{\mathrm{htl.0}}}{m^+_s / m^+_{s-1}} m^{-1/4} C_{i,s}^{(\Gamma)} B(m)^{(1-\Gamma)}$ | (A.3.11) |
| Biomass within a size range | µgC $L^{-1}$ | $B(m) = \sum_i \sum_s C_{i,s}(m_{i.s}/10^{\sigma_F/2} < m \leq m_{i.s} 10^{\sigma_F/2})$ | (A.3.12) |

**Table A.3.** Equations of the copepod model

| | Units | Equation | |
|---|---|---|---|
| Preference function for prey | - | $\phi(m_{\mathrm{py}}, m) = \exp\left[-\frac{\left(\ln\left(\frac{\beta m_{\mathrm{py}}}{m}\right)\right)^2}{2\sigma^2}\right]$ | (A.4.1) |
| Prey for copepods | µgC L$^{-1}$ | $E(m) = \sum_{i=1}^{I}\sum_{s=1}^{S} c_{\mathrm{py}}(\tau_{i,s}, m_{i,s})\Phi(m_{i,s}, m)C_{i,s} + \sum_{k=1}^{K}\Phi(m_k, m)P_k + \sum_{l=1}^{L}\Phi(m_l, m)F_l + \sum_{j=1}^{J}\Phi(m_j, m)D_j$ | (A.4.2) |
| Prey for protists | µgC L$^{-1}$ | $E_{\mathrm{u}}(m) = \sum_{k=1}^{K}\Phi(m_k, m)P_k$ | (A.4.3) |

**Table A.4.** Food available for copepods and protists

equations (ODEs, Eq. A.1.). One ODE represents the adult stage ($S$), another ODE the first stage ($s = 1$), and the rest of ODEs are the size-classes between the first stage and the adult stage. The first ODE describes the dynamics of the first copepod stage. The first terms in the right hand side is the reproduction rate of the adults, the second the growth rate within that stage (biomass accumulation), the third the maturation rate towards the next stage. The last term is the mortality rate.

Mortality in copepods originates from predation by other copepods (Eq. A.3.10), starvation (Eq. A.3.9) and a closure term representing mortality induced by higher trophic levels (Eq. A.3.11) such as fish. The mortality by higher trophic levels only applies to the copepods that are not explicitly eaten by anyone in the model, and thus mainly affects intermediate and large copepods. This mortality is a mix of density-dependence between individuals of the same size-class and individuals of similar sizes. In equation A.3.11, $\mu_{\mathrm{htl.0}}$ is the coefficient (µgC$^{1/4}$ µgC$^{-2}$ L$^{-2}$ d$^{-1}$), corrected for the number of size classes by dividing it by the ratio of the boundaries ($m_s^+/m_{s-1}^+$) of each size class. $p_{\mathrm{htl}}(m)$ is a sigmoidal function to impose the mortality only on copepods with a size larger than $m_{\mathrm{htl}} = m_{\mathrm{max}}/\beta$ ($m_{\mathrm{max}}$ is the size of the largest copepod in the community) and declines with mass $\propto m^{-1/4}$. The density dependence is also applied to whole size-ranges ($B$ Eq. A.3.12, i.e. all zooplankton biomass falling within the size range of a potential predator). The size range of $B$ is $[m/10^{\sigma_F/2} : m10^{\sigma_F/2}]$, where $\sigma_F$ is the width of the predation function of a predator, here equivalent to 1. Finally, $\Gamma$ is used to partition the density-dependence between individual stages and whole biomass size-ranges. If $\Gamma = 1$, the density dependence is imposed on each stage of each populations ($C_{i,s}$), if $\Gamma = 0$ the density-dependence is imposed on the biomass ($B$) within the size ranges. We chose $\Gamma = 0.2$.

##### A.3.3 Copepod traits

Each copepod population is characterised by the size of the adult copepod and the feeding mode. Note that juveniles have different sizes within each population. All rates follow an allometric scaling. The feeding mode of copepod can be active or passive. Active copepods include cruising copepods and feeding-current feeders, which include most calanoid copepods. Passive feeding copepods are ambush-feeders, which include some calanoids and other common copepods such as *Oithona*. Active copepods are characterised by high ingestion rates of prey and growth rates whereas they have high metabolic expenses and mortality rates. On the other hand, passive feeders will have lower ingestion rates but lower metabolism and mortality. This is implemented in the parametrization (table A.6) and predation is reduced through the preference function.

#### A.4 Protists model

The biomass in each protists size-class $P_k$ (Eq. A.2) increases with protists growth $\eta$ ($\mathrm{d}^{-1}$) and decreases with total mortality $\mu_k$ ($\mathrm{d}^{-1}$):

$$\frac{\mathrm{d}P_k}{\mathrm{d}t} = \eta_{\mathrm{u},k} P_k - \mu_{\mathrm{u},k} P_k \tag{A.6}$$

The protist model considers organisms that are mixotrophs. Protists can simultaneously do photosynthesis, take up dissolved nutrients and predate on other protists. Equations of the protists model can be found in table A.5 and A.7. Uptake of all resources (light, nitrogen, food) is performed through a type II functional response (Eq. A.5.1). Uptake rates saturate at the maximum uptake rate for resources. Food available for protists also follows the size-preference function for prey (Eq. A.4.3). Growth is the result of the most limiting resource (carbon or nitrogen), following Liebig's law of the minimum (Eq. A.5.3). Surplus of nitrogen is then leaked back to the environment (Eq. A.5.4). Protists have two sources of mortality: predation and viral lysis. Predation originates from other protists or copepods (Eq. A.5.6). Viral lysis is a density-dependent function (Eq. A.5.7).

| | Units | Equation | |
|---|---|---|---|
| **Energy gain** | | | |
| Uptake of each resource | - | $\eta_{\mathrm{X}}(m) = \overbrace{V_{\max}(m)}^{\text{Maximum uptake rate}} \dfrac{\alpha_X(m)X}{\alpha_X(m)X + g_{\max}(m)}$ | (A.5.1) |
| Respiration rate | $\mathrm{d}^{-1}$ | $\eta_{\mathrm{R}}(m) = \delta_{\mathrm{g:r}} g_{\max}(m)$ | (A.5.2) |
| Division rate | $\mathrm{d}^{-1}$ | $\nu_{\mathrm{u}}(m) = \min\left[\eta_{\mathrm{L}}(m) + \eta_{\mathrm{E}}(m) - \eta_{\mathrm{R}}(m),\ \eta_{\mathrm{N}}(m) + \eta_{\mathrm{E}}(m)\right]$ | (A.5.3) |
| Nitrogen leaks | $\mathrm{d}^{-1}$ | $\eta_{\mathrm{leaks}} = \max[0,\ \eta_{\mathrm{N}} - \eta_{\mathrm{L}} + \eta_{\mathrm{R}}]$ | (A.5.4) |
| **Mortalities** | | | |
| Total mortality | $\mathrm{d}^{-1}$ | $\mu_{\mathrm{u,tot},k} = \mu_{\mathrm{u,pr},k} + \mu_{\mathrm{u,b},k}$ | (A.5.5) |
| Predation mortality | $\mathrm{d}^{-1}$ | $\mu_{\mathrm{u,pr},k} = \sum_{i=1}^{I}\sum_{s}^{S} \dfrac{\Phi(m_k, m_{i,s})}{E_{i,s}} h m_{i,s}^{n} f_{i,s} C_{i,s} + \sum_{j=1}^{K} \dfrac{\Phi(m_k, m_j)}{E_{u,j}} \eta_{\mathrm{E},j} P_j$ | (A.5.6) |
| Viral lysis | $\mathrm{d}^{-1}$ | $\mu_{\mathrm{u,b},k} = \dfrac{\mu_{\mathrm{u,b0}}}{m_k^{+}/m_{k-1}^{+}} P_k m_k^{-1/4}$ | (A.5.7) |

**Table A.5.** Equations of the Protists model

| Symbol | Description | Units | Value | |
|---|---|---|---|---|
| | | | Active | Passive |
| $m$ | Body mass copepods | µgC $\#^{-1}$ | | |
| $m_{\mathrm{a}}$ | Body mass adult copepods | µgC $\#^{-1}$ | | |
| $m_0$ | Offspring mass | µgC $\#^{-1}$ | | |
| $z_{\mathrm{a:o}}$ | Adult to offspring mass ratio | - | 100 | |
| $m_{\mathrm{py}}$ | Body mass of a prey | µgC $\#^{-1}$ | | |
| $\beta$ | Preferred predator:prey mass ratio | - | 10000 | 100 |
| $\sigma$ | Width of prey-size function | - | 1.5 | 1 |
| $v$ | Clearance rate coefficient | L µgC$^{-3/4}$ d$^{-1}$ | 0.011 | 0.0052 |
| $q$ | Clearance rate exponent | - | $-1/4$ | |
| $h$ | Maximum ingestion rate coefficient | µgC$^{1/4}$ d$^{-1}$ | 1.37 | 0.4 |
| $n$ | Maximum ingestion rate exponent | - | $-1/4$ | |
| $k_b$ | Fraction of $h(m)$ that goes to basal metabolism | - | 0.1 | 0.1 |
| $k_{SDA}$ | Fraction of $h(m)$ that scales with feeding level | - | 0.2 | 0.2 |
| $p$ | Respiration rate exponent | - | $-1/4$ | |
| $\epsilon$ | Assimilation efficiency | - | 0.67 | |
| $\epsilon$ | Reproduction efficiency | - | 0.25 | |
| $c_{\mathrm{py}}$ | Reduced preference for passives | - | 1 | 1/5 |
| $\mu_{\mathrm{htl.0}}$ | Mortality by higher trophic levels coefficient | µgC$^{1/4}$ µgC$^{-2}$ L$^{-2}$ d$^{-1}$ | $0.003 \times h$ | |
| $Q_{\mathrm{C:N}}$ | Carbon to nitrogen ratio | - | 5.6 | |
| $\sigma_{htl}$ | Width of the prey-size function for a higher trophic level | - | 1 | |
| $\varepsilon_{\mathrm{r}}$ | Fraction of females:males and of eggs that survive | - | 0.25 | |
| $\Gamma$ | Factor scaling individual-specific v.s. size-specific mortality by higher trophic levels | - | 0.2 | |

**Table A.6.** Copepod variables and parameters. $\#$ refers to "numbers" and units of $\#^{-1}$ are "per individual". The rest of parameters and corresponding derivation are explained in Serra-Pompei et al. (2020).

#### A.5 Fecal pellets and deadfalls

Fecal pellets ($F$, Eq. A.3) are produced by copepods ($f_{\mathrm{pp}}$). Deadfalls ($D$, Eq. A.4) increases with viral lysis of protists ($\mu_{\mathrm{u,b0}}$) and a fraction $\delta$ of the mortality by higher trophic levels of copepods ($\mu_{\mathrm{htl}}$). Both fecal pellets and deadfalls are remineralized ($r$) or eaten by other copepods ($\mu_{\mathrm{p,f},l}$) without any preference function for size, as it has been shown that copepods can feed on detrital particles larger than themselves (Kiørboe, 2000; Koski et al., 2020).

| | Units | Equation | |
|---|---|---|---|
| Consumption of fecal pellets by copepods | $\mathrm{d}^{-1}$ | $\mu_{\mathrm{pr,F},l} = \sum_i^I \sum_s^S \frac{\Phi_{C\to F}}{E} h m_{is,}^n f(m_{i,s}) C_{i,s}$ | (A.7.1) |
| Consumption of deadfalls by copepods | $\mathrm{d}^{-1}$ | $\mu_{\mathrm{pr,D},l} = \sum_i^I \sum_s^S \frac{\Phi_{C\to D}}{E} h m_{is,}^n f(m_{i,s}) C_{i,s}$ | (A.7.2) |

**Table A.7.** Consumption of Fecal pellets and deadfalls by copepods. $\Phi_{C\to F}$ refers to the preference of copepods for fecal pellets (the same for deadfalls).

#### A.6 Nitrogen

Dissolved nitrogen ($N$, Eq. A.5) is taken up by protists $\eta_N$. Its concentration increases through bacterial remineralization $r$, leakage from protists $\eta_{\mathrm{leaks}}$ and excretion from copepods $\eta_{\mathrm{DON}}$. Higher trophic levels, not explicitly considered in the model, respire carbon and excrete nitrogen back to the environment. Thus, we assume that a fraction $1 - \delta$ of the losses to higher trophic levels $\mu_{\mathrm{htl}}$ is remineralized. Inputs and mixing of nitrogen are taken care by the transport matrix model.

#### A.7 Temperature dependencies

Temperature effects are implemented as factors on the relevant parameters of copepods and protists. We use $Q_{10}$ factors to model the effects of temperature on each corresponding parameter, which for a given rate $R$ has the following effect:

$$R = R_{\text{ref}} Q_{10}^{(T - T_{\text{ref}})/10}, \tag{A.7}$$

where $T$ is temperature, $T_{\text{ref}}$ the reference temperature and $R_{\text{ref}}$ the rate at the reference temperature. Temperature-dependent parameters are listed in table A.8, alongside their corresponding $Q_{10}$.

| Symbol | Description | $Q_{10}$ |
|---|---|---|
| $h$ | Maximum ingestion rate copepods | 2 |
| $v$ | Clearance rate coefficient | 1.5 |
| $\alpha_N$ | Affinity for nitrogen | 1.5 |
| $\alpha_L$ | Affinity for light | 1 |
| $\alpha_F$ | Affinity for food | 1.5 |
| $V_{\max,X}$ | Maximum uptake rates of resources protists | 2 |
| $V_{\max,X}$ | Respiration rate protists | 2 |
| $r$ | Remineralization rate | 2 |
| $\mu_{\text{htl}}$ | coefficient of higher trophic level mortality | 2 |

**Table A.8.** Temperature dependencies affecting different rates.

#### A.8 Physical model

The NUM framework is embedded in a physical model of the global ocean. Physics are simulated by the transport matrix method (Khatiwala, Visbeck, and Cane, 2005; Khatiwala, 2007). We use a coarse resolution (2.8° $\times$ 2.8°, 15 vertical levels), monthly-averaged transport matrix from the MIT gcm (`http://kelvin.earth.ox.ac.uk/spk/Research/TMM/TransportMatrixConfigs`, as used in Dutkiewicz, Follows, and Parekh (2005)). The coarse resolution results in the euphotic zone being resolved only by the two first layers of the transport matrix (at most up to the third layer in clear waters). The temperature forcing is also monthly averaged and was provided in the transport matrix itself. Irradiance at the ocean surface was taken from the Ocean Productivity site. The data was afterwards interpolated to fit the grid of the transport matrix. The irradiance experienced by primary producers in each layer is as follows: surface irradiance is $I_0$ and the incoming irradiance in the upper boundary ($z_{i,up}$) of each depth-layer $z_i$ is:

$$I_{z,up} = I_0 e^{-k_{tot} z_{i,up}}, \tag{A.8}$$

where $k_{tot}$ is the attenuation coefficient, which depends on the attenuation coefficient of seawater $k_w$ ($m^{-1}$) and on the attenuation by planktonic particles $k_p$ ($m^2$ $mgC^{-1}$):

$$k_{tot} = k_w + k_p \int_0^{z_i} P \mathrm{d}z. \tag{A.9}$$

The realised light experienced by plankton in each depth layer is the mean of the light level in the depth bin:

$$I_z = \frac{I_{z,up}}{k_{tot} \Delta_{z,i}} (1 - e^{-k_{tot} \Delta_{z,i}}), \tag{A.10}$$

where $\Delta_{z,i}$ is the depth range in each grid of the transport matrix.

#### A.9 Carbon export

The sinking rate of fecal pellets and deadfalls is implemented as a simple first order upwind scheme in the transport matrix. Hence, the rate of change of a state variable $X$ due to sinking $v_{\text{sink}}$ in each depth bin ($z$) is:

$$\frac{\mathrm{d}X_z}{\mathrm{d}t} = \frac{v_{\text{sink}}}{\Delta_z} (X_{z-1} - X_z); \tag{A.11}$$

where $X$ denotes deadfalls or fecal pellets particles concentration in a bin ($z$), $v_{\text{sink}}$ the sinking rate (in m $d^{-1}$) associated to the particle and $\Delta_z$ the depth-range of the bin (in m). Therefore this sinking term affects the ODEs from $F$ and $D$ (Eq. A.3 and A.4 respectively).

The sinking rate attributed to each particle depends on the particle size $m$ within each size-class (Fig. A.10). Fecal pellets sinking rates are relatively well constrained, and we use the data from

Small, Fowler, and Ünlü (1979). Deadfalls sinking rates are highly uncertain since the particles might change size as they sink by coagulation, fragmentation or changes in density. We do not model these processes and assume a sinking rate that is only weakly dependent on particles size, consistent with observations (Alldredge and Gotschalk, 1988).

##### A.10 Carbon export efficiency (pe-ratio)

The pe-ratio is defined as the fraction of depth-integrated NPP exported as sinking particles at a given depth horizon. The depth horizons used in this study are 120 m and 1080 m, which are the bottom of the second and the seventh layer of the transport matrix respectively. We consider the annual and seasonal pe-ratio. The annual pe-ratio is the ratio between NPP and export flux integrated over a year. The seasonal pe-ratio is on a shorter time scale, and is the daily particle flux divided by NPP, the latter averaged over a given time frame. We used 3 NPP-averaged time frames to see how it might affect the value of pe-ratio: (i) the instantaneous NPP (NPP at the same time of export), (ii) 15 days averaged, and (iii) 30 days averaged.

#### B Trophic level calculation

The trophic level of each size-class of protists and copepods was calculated iteratively (Ward and Follows, 2016). Given a total number of protists and copepod size classes $N$, the trophic level $T_i$ of a compartment $i$ is:

$$T_i = 1 + \sum_{j=1}^{N} T_j p_{ij}(1 - p_{\text{auto}}) \tag{B.12}$$

Where $T_j$ is the trophic level of each prey $j$, $p_{ij}$ is the fraction that the prey $j$ contributes to the total ingestion of predator $i$, and $p_{\text{auto}}$ is the autotrophic fraction of mixotrophs (i.e. $p_{\text{auto}} = 1$ if a protist growth only from photosynthesis, and is 0 if it is fully heterotrophic. Therefore, pure autotrophs have a trophic level of 1, and heterotrophs a level higher than 1. Since we allow for mixotrophic feeding, there can be intermediate trophic levels. For instance, a mixotroph that gains half of its energy from photosynthesis and the other half from preying on an autotroph with trophic level 1 would have a trophic level equal to 1.5. This approach can also result in some mixotrophs having a trophic levels lower than their prey. This is because, despite them feeding slightly on that prey, they are mainly autotrophs, and therefore the autotrophic scaling reduces their trophic level. Copepods have a $p_{\text{auto}} = 0$).

To calculate $p_{\text{auto}} = 1$ we first consider which is the limiting factor: carbon or nitrogen. If carbon is limiting, the autotrophic fractions is:

$$p_{\text{auto}} = \frac{J_L}{J_L + J_F} \tag{B.13}$$

If nitrogen is limiting, the autotrophic fractions is:

$$p_{\text{auto}} = \frac{J_N}{J_N + J_F} \tag{B.14}$$

#### C Net primary production

To calculate net primary production, we need to consider whether the respiration of mixotrophs originated from photosynthesis or from ingestion of prey. In addition, net carbon fixed is only the one used for growth. We also do not allow to have negative NPP (i.e. when overall respiration is higher than the photosynthetic rate). Hence NPP in the model is calculated as:

$$NPP = \sum(\max[0, J_L - J_R p_{\text{auto}} - \max(0, J_L - J_R - J_N)]P); \tag{C.15}$$

The first "max" statement prevents from having negative NPP (i.e. when respiration is higher than the photosynthetic rate). The second "max" statement accounts for the effects of nitrogen limitation. Following our assumptions, the cell will only fix the carbon that is allowed by the available nitrogen in the cell following the stoichiometric ratio. Hence, excess carbon "fixed" is not accounted as NPP as it does not go into cell growth (in nature it would go into DOC, but we do not have this compartment here).

### D Supplementary figures

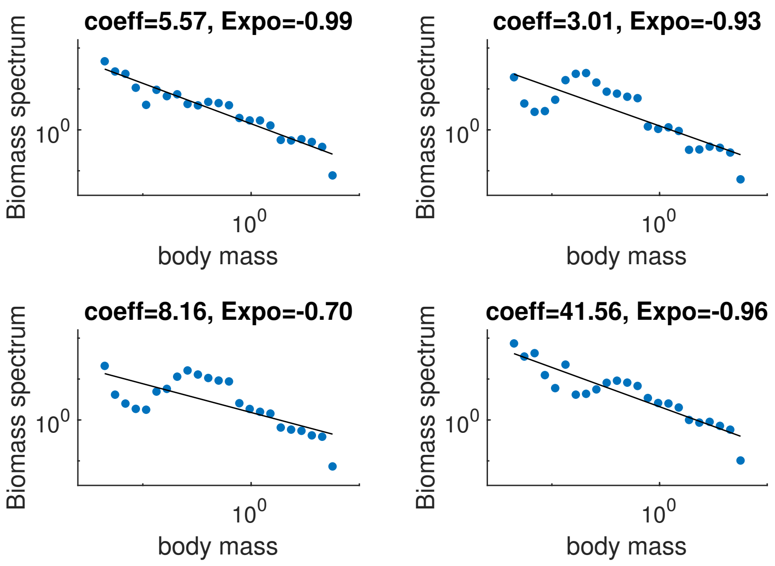

**Fig. D.1.** Example of modelled binned community spectra (dots) used to obtain the exponent ($\lambda$) and the coefficient ($\kappa$) used in the main text. The linear regression was done by using a least squares method on: $\log_{10}(B_{spec}) = \lambda \log_{10}(m) + \log_{10}(\kappa)$.

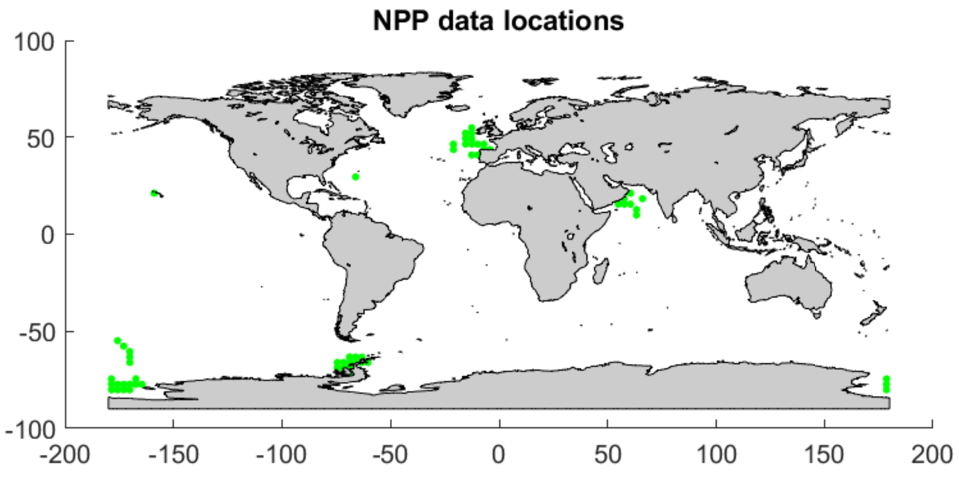

**Fig. D.2.** Location of the NPP data used to validate the model. Data compiled in Saba et al. (2011).

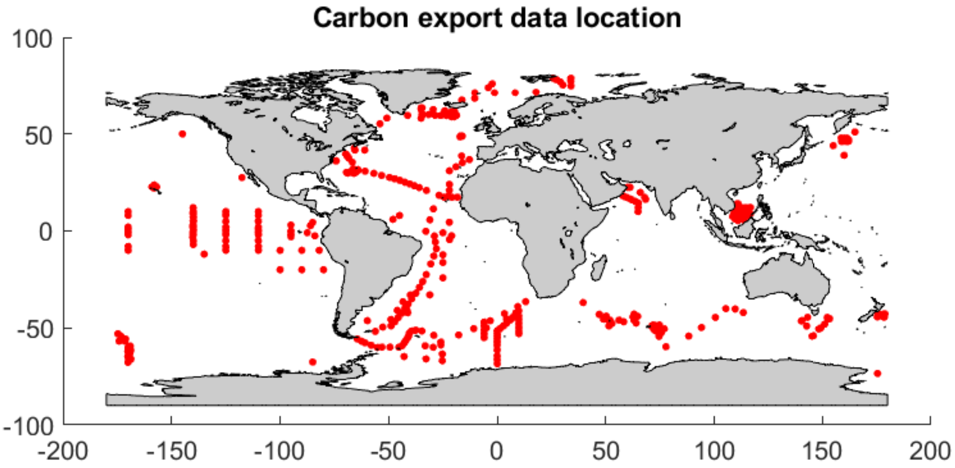

**Fig. D.3.** Location of the carbon export data used to validate the model. Data taken from Henson, Le Moigne, and Giering (2019) and Le Moigne et al. (2013).
